# A senescence-like state is beneficial for ovarian cancer treatment

**DOI:** 10.1101/2023.09.30.560300

**Authors:** Michael Skulimowski, Llilians Calvo, Shuofei Cheng, Isabelle Clément, Lise Portelance, Yu Zhan, Euridice Carmona, Julie Lafontaine, Manon de Ladurantaye, Kurosh Rahimi, Diane Provencher, Anne-Marie Mes-Masson, Francis Rodier

## Abstract

High-grade serous ovarian carcinoma (HGSOC) commonly responds to initial therapy, but this response is rarely durable. Understanding cell fate decisions taken by HGSOC cells in response to treatment could guide new therapeutic opportunities. Here we find that tissue-derived primary HGSOC epithelial cultures reflecting the original disease primarily undergo therapy-induced senescence in response to DNA damage and first-line carboplatin/paclitaxel chemotherapy. Unlike previous observations using cell lines, primary HGSOC cell TIS displays a stable senescence proliferation arrest and p16INK4A expression, and is accompanied by persistent DNA damage, an inflammatory secretome, and senolytic sensitivity, suggesting new avenues for selective pharmacological manipulation of these cells. Whether cell senescence induced by cancer therapy is beneficial or detrimental to clinical outcomes remains unknown. Single cell comparison of pre- and post-chemotherapy patient HGSOC tissue samples revealed changes in physio-pathological senescence biomarkers supporting a post-treatment senescence-like state. Importantly, patients with stronger senescence signatures post-chemotherapy displayed better 5-year survival suggesting that senescence accounts, at least in part, for beneficial cellular responses to treatment. Given that ovarian cancer epithelial cells almost universally retain the capacity to undergo senescence and that senescence appears beneficial in this context, HGSOC senescence-centric therapeutic avenues should be further explored.

**Author Summary:** Whether cancer therapy-induced cell fate decisions like senescence are good or bad for patient treatment outcome is unknown. We find that ovarian cancer cells almost universally retain senescence competence and that senescence in treated cancer tissues correlate with good clinical outcomes. This reveals that senescence is a relevant drug target in ovarian cancer.

## INTRODUCTION

Ovarian cancer remains the most lethal gynecological malignancy in the United States and Canada. Among the different subtypes of the disease, high-grade serous ovarian cancer (HGSOC) is by far the most common, making up roughly two thirds of cases (1). Currently, initial treatment consists of a cytoreductive surgery accompanied by six cycles of a platinum- and taxane-based combination chemotherapeutic regimen (2). Although primary response rates are high, toxicities associated with multiple rounds of chemotherapy and eventual cancer cell drug resistance limit 5-year overall survival to around 40% (3). Nevertheless, the high initial response rates suggest that a greater understanding of the molecular basis of the cellular responses involved could provide avenues to improve therapeutic approaches.

Cellular responses to stress, such as DNA damage induced by radio- or chemotherapy, are termed cell fate decisions. These include repair or bypass of the damage, cell death (programmed or catastrophic), and cellular senescence (permanent growth arrest) (4, 5). Senescent cells are characterized by several senescence-associated (SA) hallmarks that contribute to their biological functions. Among these, the most prominent are the SA proliferation arrest (SAPA), the micro-environmentally active SA secretory phenotype (SASP), and enhanced survival via SA apoptosis resistance (SAAR) (5–9). The SAPA functions as a direct cancer suppression mechanism and is largely mediated by two key tumor suppressor pathways, p53/p21^WAF1^ and p16^INK4A^/Rb, although many other pathways contribute a large degree of redundancy in this program (7, 10).

The stability of the SAPA is maintained to a great extent through persistent DNA damage response (DDR) signaling and via SA chromatin remodeling associated with the expression of the cyclin-dependent kinase inhibitor (CDKi) p16^INK4A^(5–7). Though SA chromatin remodeling remains poorly understood, the downregulation of nuclear laminB1 seems to facilitate this process, and is also a senescence hallmark (11, 12). The SASP is composed of growth factors, cytokines, chemokines, extracellular proteases, and extracellular matrix proteins that mediate senescent cell functions in tissue repair and interactions with other cells including immune cells (5, 7, 9, 13). Finally, SAAR reflects augmented anti-apoptotic mechanisms including the activity of antiapoptotic Bcl-2 family members. This results in senescent cells being uniquely sensitive to Bcl2/Bcl-XL inhibitors, which are among founding members of a new class of drugs termed “senolytics” (14–17). In the context of carcinogenesis, senescence is undoubtedly beneficial in its role as a barrier to malignant transformation in pre-neoplastic lesions (5–7). Alternatively, therapy-induced senescence (TIS) in damaged cancer cells is advantageous by preventing tumor cell growth (10, 18, 19). However, it remains unclear whether TIS cancer cells have stably stopped proliferation, fueling suggestions that these cells may represent dormant reservoirs contributing to treatment resistance (10, 20). Similarly, alterations to the tissue microenvironment caused by TIS-SASP must be evaluated contextually. For example, in a mouse model of liver cancer, the microenvironment created by the presence of senescent cells was shown to promote the immune clearance of damaged cancer cells, suggesting that senescence could enhance the therapeutic response (21). In contrast, TIS was demonstrated to promote treatment resistance in a mouse lymphoma model by creating DNA damage-induced survival niches for residual cancer cells (22). Currently, the role of TIS as an overall beneficial or detrimental process during human cancer treatment remains largely unknown.

A few studies have begun to shed light on the relevance of senescence in ovarian cancer. For example, evidence of persistent DDR signaling, often associated with senescence, have been reported in human hyperplastic fallopian tubes (23). This suggests that senescence acts as a tumor suppression barrier in this context to prevent the progression from pre-neoplastic lesions to HGSOC, perhaps applying a selective pressure responsible for the extremely high prevalence of p53 mutations in this disease (24, 25). After malignant transformation, the induction of senescence is reported in immortalized cell lines derived from various ovarian epithelial cancer subtypes. Briefly ovarian cancer cell lines can be forced into senescence via drug exposure including poly(ADP-ribose) polymerase 1 inhibitors (PARPi) (18, 26, 27), growth factor overexpression (28), steroid exposure (29), and miRNA overexpression (30, 31). In HGSOC specifically, IDH1 suppression in cell lines triggers senescence (32), and PARPi trigger a reversible senescence-like state that can be opportunistically targeted using senescence-centric approaches (26). Despite these advances, many studies suggest that immortalized ovarian cancer cell lines may not reflect the full spectrum of human disease due to clonal derivation and low spontaneous immortalization rates, which is well established for HGSOC (33–35).

Here, using low passage human HGSOC primary cultures, patient tissues, and their associated clinical data, we find that culture stress and chemotherapy-induced DNA damage almost universally trigger senescence in HGSOC cells. HGSOC cells undergoing TIS are found to harbor senescence hallmarks like persistent DDR signaling and to express a SASP. Importantly, senescent HGSOC cells are found to be selectively sensitive ABT-263, a senolytic agent that inhibits Bcl-2 and Bcl-xL (14, 26). Comparison of pre- and post-chemotherapy patient tissues reveals changes in biomarkers suggestive of senescence after treatment. Lastly, correlation of the expression of senescence biomarkers in post-chemotherapy tissues with clinical data shows TIS to be associated with more favorable clinical outcomes. Overall our findings suggest that TIS is potentially beneficial in HGSOC and may be a therapeutic target for the improvement of current.

## RESULTS

### HGSOC primary cultures undergo culture stress-induced senescence

To define a human disease model adequate for the study of HGSOC therapy-induced cell fate decisions (TICFD), we first retrospectively reviewed data from the long-standing Centre hospitalier de l’Université de Montréal (CHUM) ovarian cancer tissue bank, a resource that systematically attempted to derive cell lines from serial ovarian cancer primary cultures and which, over the years, has generated more than 30 ovarian cancer cell lines. Primary culture data reveals that the vast majority of hundreds of primary HGSOC cultures did not spontaneously immortalize into cancer cell lines. Instead, following limited proliferation in culture for a few passages, growth slowed and eventually stopped, and cells adopted a flattened enlarged morphology without evidence of extensive cell death, suggestive of senescence. For example, out of 86 primary HGSC cultures in the year 2000, only one culture generated an immortal cell line, while the remaining cultures ceased proliferation after an average of 4.6 passages (Fig. 1a-b). Following optimization of culture conditions (years 2006-2007), 15 HGSOC cell lines were established out of a total of 187 primary cultures, yielding a success rate of 8.02% consistent with other observations (33–35) and reinforcing the idea that spontaneous immortalization of HGSOC primary cells is a relatively rare event.

**Figure 1:**
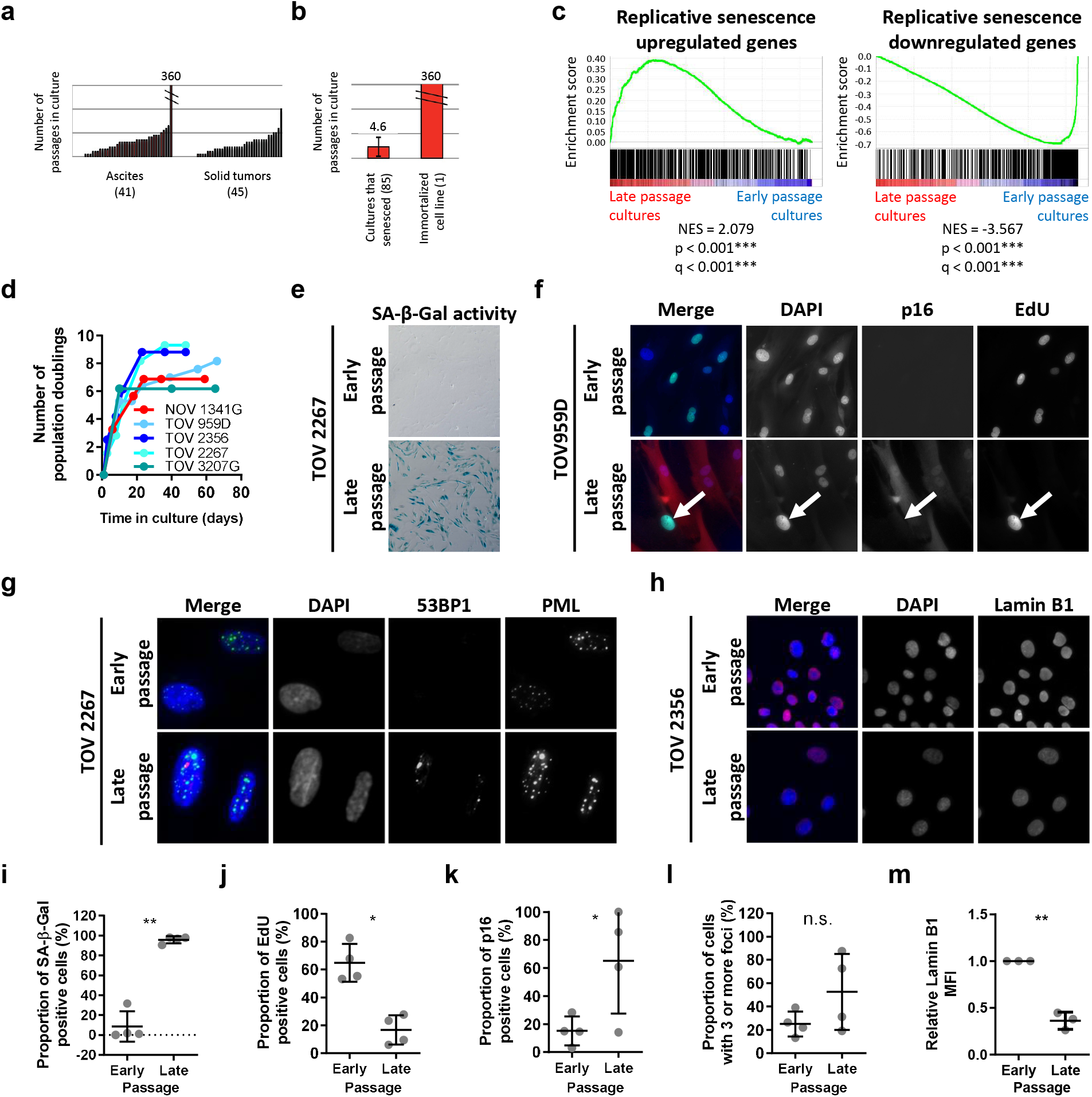
HGSOC primary cultures undergo culture stress-induced senescence. (**a**) The maximum number of passages before proliferation arrest of 86 independent HGSOC primary cultures collected during the year 2000. (**b**) The average maximal number of passages for growth-arrested cultures (85/86) or spontaneously immortalized cultures (1/86). (**c**) Gene set enrichment analysis (GSEA) plots of differentially expressed genes in late and early passage HGSOC cultures for genes found to be upregulated (left) and downregulated (right) in the normal human fibroblast replicative senescence gene expression signature (Hernandez-Segura et al., 2017). (**d**) Proliferation curves of normal epithelial ovarian primary cells (NOV1341G) and primary HGSOC cells (TOV959D, TOV2356, TOV2267, and TOV3207G) serially passaged until proliferating arrest. Data is reported as population doublings (PD) over time. (**e**) Early- and late-passage HGSOC (TOV959D, TOV1294G, TOV2267, and TOV2356) primary cultures were stained for SA-β-Gal activity (representative TOV2267 images shown). (**f**) Early- and late-passage HGSOC primary cultures (TOV959D, TOV1294G, TOV2267, and TOV2356) were stained for p16INK4A (red) and EdU (24-hour pulse; green) using immunofluorescence. Nuclei are counterstained with DAPI (blue) (white arrow highlights an EdU-positive, p16-negative cell) (representative TOV959D images shown). (**g**) Early- and late-passage HGSOC primary cultures (TOV959D, TOV1294G, TOV2267, and TOV2356) were stained for 53BP1 and PML. DNA-SCARS are double-positive for both 53BP1 and PML nuclear bodies (representative TOV2267 images shown). (**h**) Early- and late-passage HGSOC primary cultures (TOV1294G, TOV2267, and TOV2356) were stained for lamin B1 (red) and nuclei counterstained with DAPI (blue) (representative TOV2356 images shown). (**i**) Quantification of the percentage of SA-β-Gal positive cells from (e). (**j**) Quantification of the percentage of EdU-positive nuclei from (f). (**k**) Quantification of p16INK4A-positive cells from (f). (**l**) Quantification of cells with more than three 53BP1 DNA damage foci from (h). (**m**) Quantification of the mean fluorescent intensity (MFI) of peripheral nuclear lamin B1 from (h) normalized to the control of each primary culture. Peripheral nuclear lamin B1 MFI was quantified within a 6 μm wide area along the perimeter of the nuclear area. Statistical significance was calculated using the paired t test for all panels except panel b, where the Kolmogorov-Smirnov statistic was used. n.s., not significant; * p < 0.05; ** p < 0.01.

We further explored this apparent senescence response using Affymetrix gene expression analysis profiles from 42 independent HGSOC primary cultures collected at various passages within the culture lifespan (from early to late passages). Individual cultures were first assigned a proliferation status using unsupervised clustering for E2F-response gene activity (Fig. S1). Proliferating early passage cells (≤50% lifespan completed) were then compared to slow-proliferating late passage cells (≥75% lifespan completed) through gene set enrichment analysis (GSEA). We found multiple senescent signature gene sets to be significantly enriched accordingly in each group of cultures (15/22 gene sets; Fig. S1d). Importantly, significant enrichment for genes upregulated in replicative senescence was found in the slow-proliferating late passage cultures, and, conversely, significant enrichment for genes downregulated in replicative senescence was found in the proliferating early-passage cells (Fig. 1c).

To further confirm key senescence hallmarks directly, we serially passaged four HGSOC primary cultures (TOV959D, TOV2356, TOV2267, TOV3207G). Similar to normal ovarian surface epithelial cells (NOV1341G) and as expected during culture stress-induced senescence (36, 37), HGSOC primary cultures gradually ceased proliferation (Fig 1d). When compared to early passage cells from their matched parental culture, late passage normal and cancer cells displayed increased senescence-associated beta-galactosidase activity (SA-β-Gal) (Fig. 1e, i, Fig. S2a) and decreased DNA synthesis (Fig. S2b-c). Tumor suppressor genes driving senescence are often mutated in cancer (10, 38). Indeed, the well-known tumor suppressor *TP53*, a master mediator of damage-induced senescence, is mutated in >95% cases of HGSOC (24, 39, 40) (Fig. S2d). However, a quick analysis of The Cancer Genome Atlas (TCGA) large ovarian cystadenocarcinoma cohorts (currently 617 combined cases) revealed that p16^INK4A^ (gene *CDKN2A)*, another important mediator of SAPA and a senescence biomarker (5–7), is mutated in less than 5 % of HGSOC cases (Fig. S2d). We thus performed an immunostaining for p16^INK4A^ concomitantly with a DNA synthesis assay on early and late passage HGSOC cells. Compared to early passage cells, a significantly higher proportion of late passage cells were positive for p16^INK4A^ and, accordingly, a significantly higher proportion of late passage cells were negative for DNA synthesis. Importantly, individual cells that stained positive for p16^INK4A^ were DNA synthesis negative, corroborating the integrity of this SAPA pathway in HGSOC primary cells (Fig. 1f, j-k). We then explored the SA accumulation of permanent DNA damage foci termed DNA-SCARS (DNA segments with chromatin alterations reinforcing senescence) (41). To this end, we co-stained for 53BP1 and PML nuclear bodies using immunofluorescence (IF), the colocalization of which has been shown to mark DNA-SCARS. Although DNA-SCARS were detected via colocalization, we quantified a non-significant increase in the proportion of cells that harbored three or more 53BP1 foci (Fig. 1g-l), perhaps owing to high levels of basal DNA damage associated with HGSOC (42, 43). Lastly, we explored lamin B1 levels, shown to be downregulated in senescent cells (11, 12), and found that late passage HGSOC cells strongly downregulated nuclear lamin B1 as compared to their early passage counterparts (Fig. 1h-m, Fig. S2e). Taken together, these data suggest that most if not all HGSOC primary cells retain the ability to undergo culture stress-induced senescence, and this senescence is associated with SAPA, SA-β-Gal, nuclear lamina alterations, the expression of p16^INK4A^, and the accumulation of additional DNA damage.

### HGSOC primary cultures undergo TIS in response to DNA damage and chemotherapy

We next tested whether HGSOC primary cells undergo TIS in response to DNA damage induced by therapy. Five different cultures of primary cells were treated with either 10 Gy ionizing radiation (IR), in order to induce senescence-associated DNA double-stranded breaks, or a combination of 10 μM carboplatin and 30 nM paclitaxel for a limited period of 12 hours (CP) to mimic one cycle of chemotherapy in patients (41, 44, 45). We then evaluated senescence hallmarks 8 days after the beginning of treatment. For both IR and CP, when compared to their controls, treated cells stained positive for SA-β-Gal (Fig. 2a,e), were largely positive for p16^INK4A^, negative for DNA synthesis (Fig. 2b, f-g), exhibited increased DNA-SCARS (Fig. 2c, h, Fig. S3a), and expressed lower levels of nuclear lamin B1 (Fig. 2d, i, Fig. S3b). Senescence induced by DNA damage has been well described to be associated with a SASP (5, 7, 9, 13).

**Figure 2:**
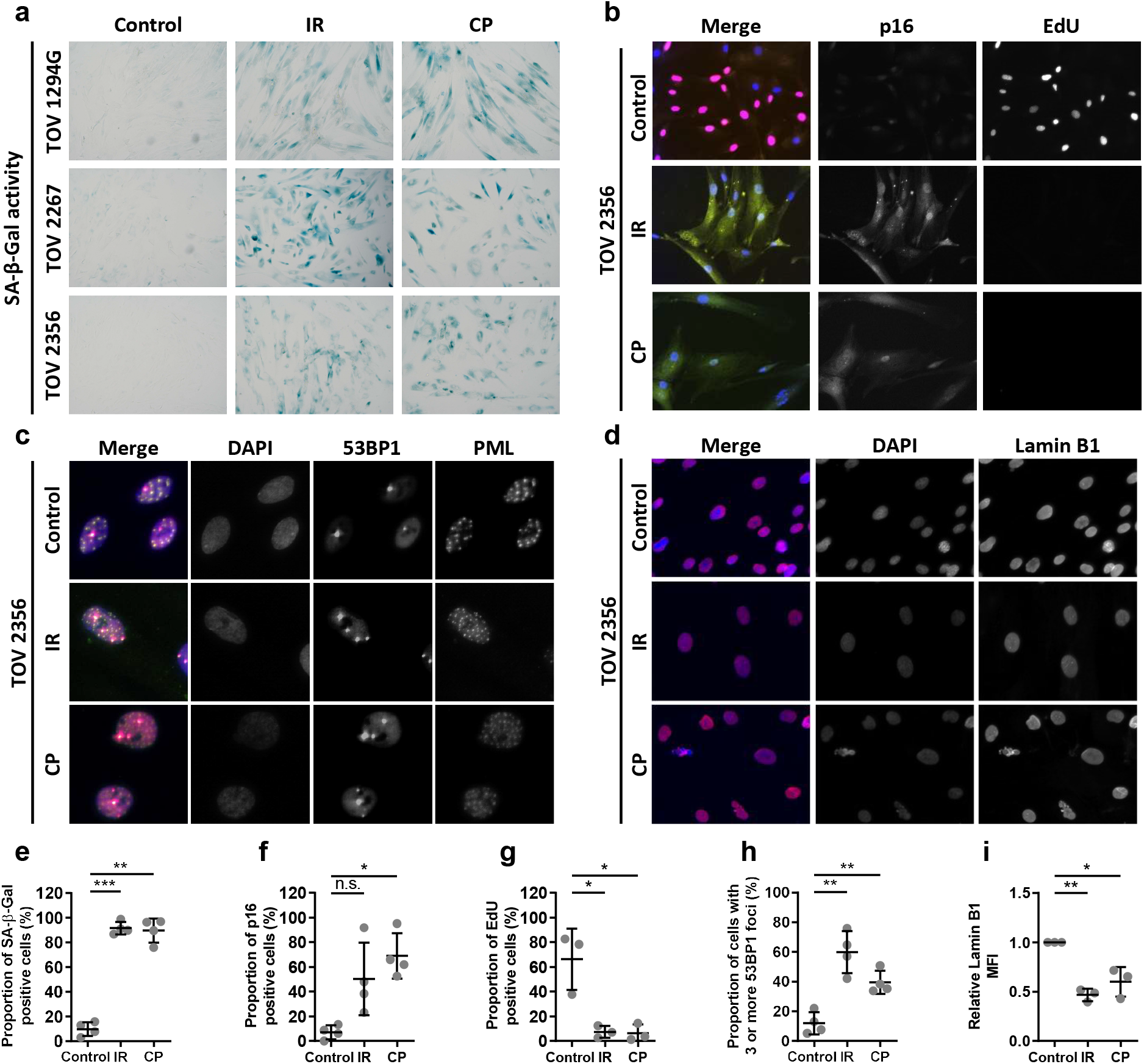
HGSOC primary cultures undergo TIS in response to DNA damage and chemotherapy. (**a**) HGSOC primary cultures were stained for SA-β-gal 8 days following treatment with 10 Gy ionizing radiation (IR) or a combination of 10 μM carboplatin and 30 nM paclitaxel for 12 hours (CP). (**b**) HGSOC primary cells were labelled with EdU (24-hour pulse; red) and stained for p16INK4A (green) 8 days after IR or CP treatment (representative TOV2356 images shown). (**c**) HGSOC primary cultures were stained for 53BP1 (red) and PML nuclear bodies (green) 8 days after IR or CP treatment. Colocalization of 53BP1 and PML (yellow in the merge panel on the left) reveals the presence of DNA-SCARS. (**d**) HGSOC primary cultures were stained for lamin B1 (red) 8 days after IR or CP treatment. (**e**) Quantification of the percentage of SA-β-gal-positive cells after IR or CP treatment from (a) in TOV 513, TOV1294G, TOV 2267, and TOV 2356. (**f**) Quantification of p16INK4A-positive cells in TOV513, TOV1294G, TOV2267, and TOV2356 cultures 8 days after treatment from (b). (**f**) Quantification of EdU-positive cells in TOV1294G, TOV2267, and TOV2356 cultures 8 days after treatment from (b). (**h**) Quantification of cells with more than three 53BP1 foci in TOV513, TOV1294G, TOV2267, and TOV2356 cultures in either untreated conditions or after IR or CP treatment from (c). (**i**) Quantification of peripheral nuclear lamin B1 MFI normalized to control in TOV1294G, TOV2267, and TOV2356 after IR or CP treatment from (d). Peripheral nuclear lamin B1 MFI was quantified in a 6 μm wide area along the perimeter of the nuclear area. Statistical significance was calculated using the paired t test. * p < 0.05; ** p < 0.01; *** p < 0.001.

Thus, we probed the secretome in two HGSOC primary cultures (TOV2267, TOV2356) using a multiplex immunoassay for a variety of SASP factors. Eight days after treatments cells displayed a strong upregulation of multiple SASP factors with specific factors differing slightly between treatments and individual cultures (Fig. 3a-b), consistent with previously observed context-dependent variations in SASP (8, 46). Therefore, HGSOC primary cells seem to undergo TIS in response to DNA damaging therapies, including treatment with a standard chemotherapeutic regimen.

**Figure 3:**
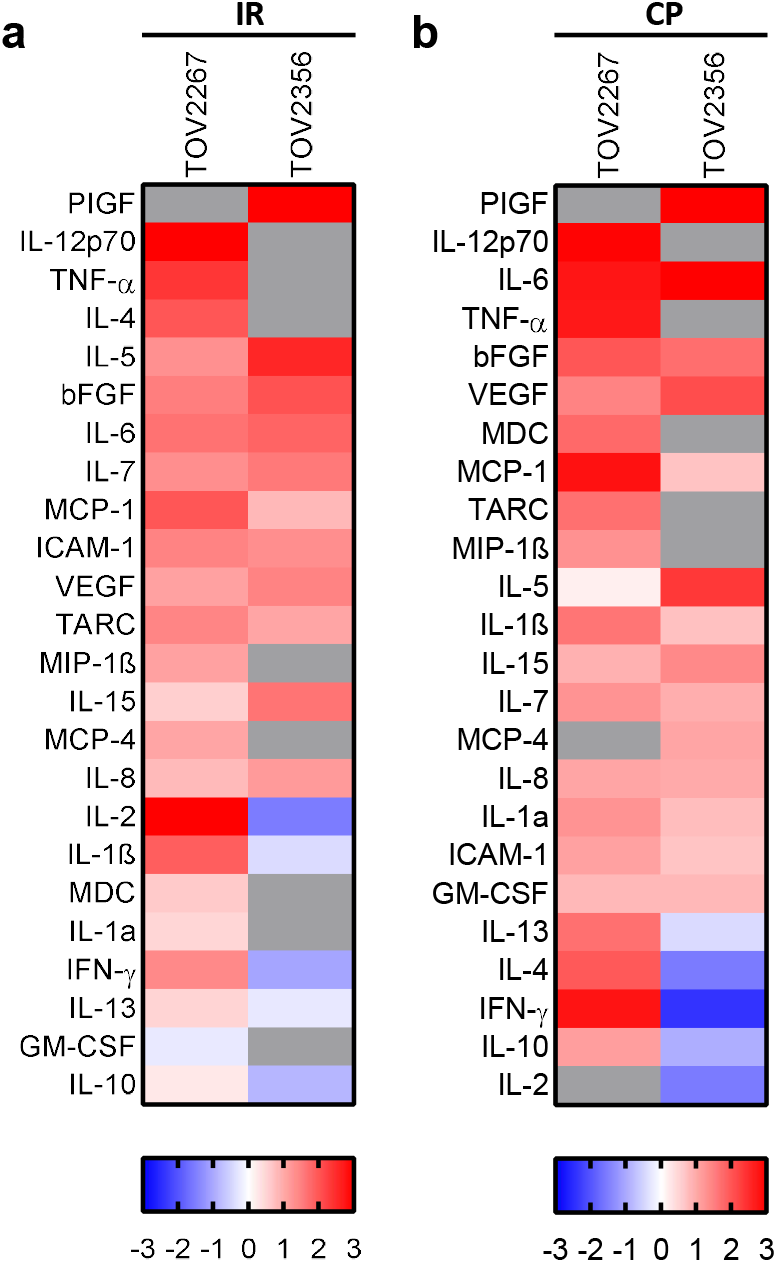
HGSOC primary cells trigger SASP secretome in response to therapy. (**a-b**) Multiplex-based SASP profiling was performed on conditioned media from TOV2267 and TOV2356 HGSOC primary cultures treated (or not) with either 10 Gy IR (**a**) or carboplatin (10 μM) and paclitaxel (30 nM) (CP) for 12 hours (**b**). The conditioned medium was prepared 8 days following treatment (serum-free OSE medium for 16 hours). The heat-map colors indicate the relative fold change on a log2 scale in protein levels (red for increased levels, blue for decreased levels relative to the control signal for each culture). Soluble factors with the highest fold changes are shown here. Factors are ordered, from top to bottom, according to decreasing fold change. Gray: not detectable or not quantifiable; IR: ionizing radiation; CP: carboplatin and paclitaxel.

### HGSOC TIS renders cells sensitive to ABT-263

SAAR is a pharmacologically targetable senescence hallmark that represents an opportunity for age-associated diseases treatment (14–17). Similarly, senescent cancer cell lines can be selectively re-directed to death using senolytic compounds like ABT-263 that inhibit the anti-apoptotic proteins Bcl-2 and Bcl-xL (20, 26, 47, 48). Given the unpredictable impact of the SASP on tissue microenvironments and the possibility that HGSOC cell SAPA is not permanently stable, we sought to determine whether ABT-263 could be used to clear senescent HGSOC cells. We first treated non senescent TOV2267 cells with increasing concentrations of ABT-263 for 72 hours and assessed relative viability (Fig. 4a-c), revealing that non senescent HGSOC cells are not sensitive to ABT-263 concentrations of up to 10 μM, much above the predicted active concentration of the compound (∼0.1-1 µM) (49). Unlike their pre-senescent counterparts, similarly treated senescent TOV2267 cells (ABT-263 treatment from day 7 post-IR) showed decreased relative viability with concentrations of ABT-263 as low as 0.1 μM, with cells detaching and floating after ABT treatment (Fig. 4a-b, d). Similarly, non senescent TOV1294G and TOV513 cells were insensitive to ABT-263 at lower doses (Fig. 4e, Fig. S4a-b), but IR- or CP-treated senescent TOV1294G and TOV513 HGSOC cells were selectively and preferentially eliminated by a 5 μM ABT-263 treatment (Fig. 4a, f-g). Thus, in addition to developing a TIS phenotype in response to therapy, HGSOC primary cells develop a sensitivity to ABT-263, suggesting a senolytic pharmacological strategy that could enhance cancer cell elimination during current therapies.

**Figure 4:**
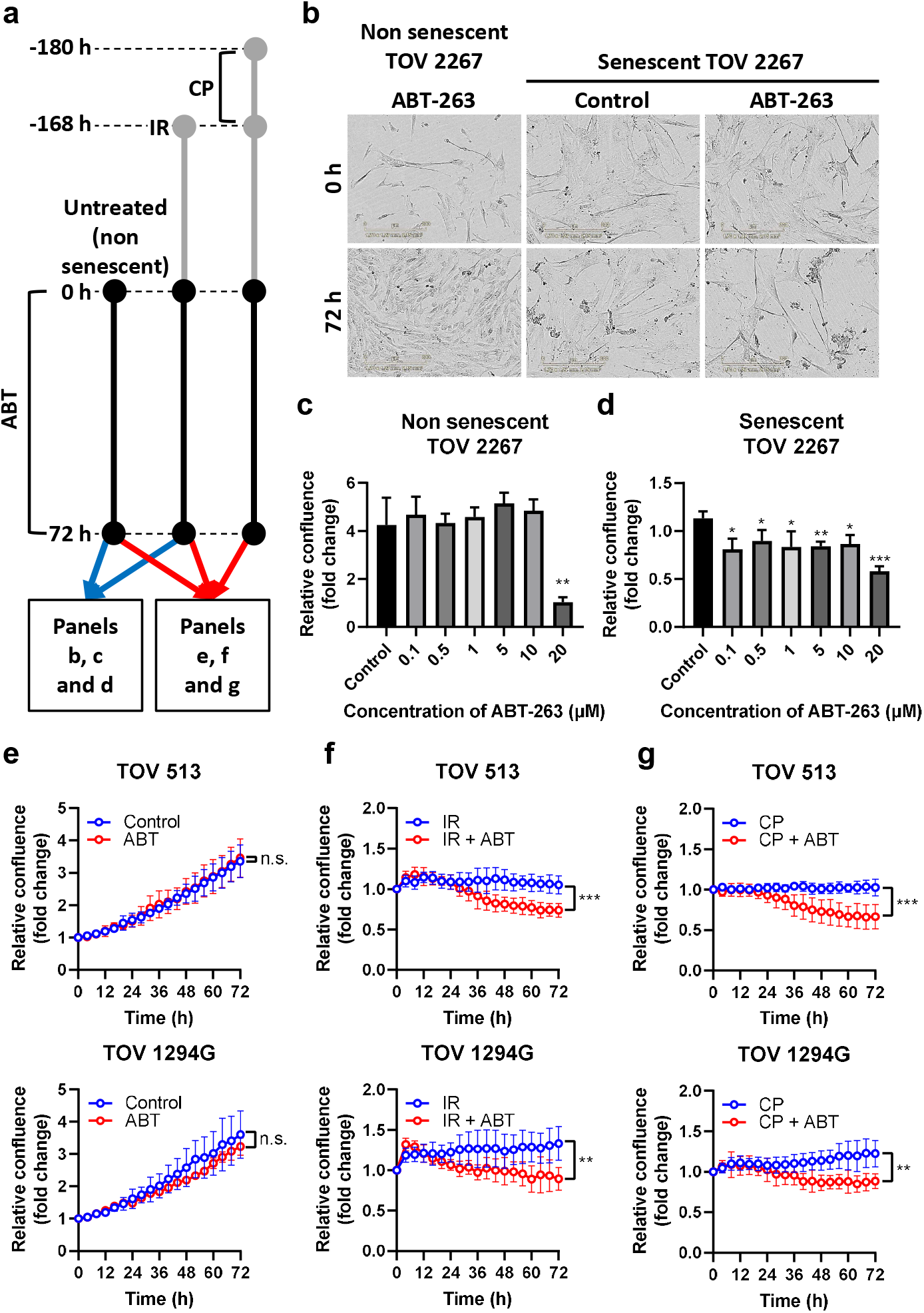
HGSOC TIS renders cells sensitive to senolytic clearance via ABT-263. (**a**) Experimental design for the present figure. Cells were seaded 24 hours prior to treatment. Non senescent cells went untreated prior to ABT-263 treatment, whereas senescent cells underwent 10 Gy IR or a 12-hour pulse of 10 uM carboplatin and 30n nM paclitaxel (CP) 7 days prior to treatment with ABT-263. Cells were then treated with varying concentrations of ABT-263 for a period of 72 hours. Confluence of the cells in condition was measured at the beginning of the ABT-263 treatment and every 4 hours thereafter or 72 hours after the initiation of ABT-263 treatment. (**b**) Non senescent or senescent (7 days after 10 Gy IR) TOV 2267 cells were treated with 5 μM of ABT-263 for 72 hours. (**c-d**) Quantification of the relative confluence of nonsenescent (**c**) and senescent (**d**) TOV2267 cells when treated with different concentrations of ABT-263 for 72 hours. Data is represented as mean of the ratio of the confluence after 72 hours of treatment over the confluence at the beginning of treatment. (**e-g**) Quantification of the relative confluence over time of non senescent (**e**) or senescent (**f**, 7 days after IR treatment; **g**, 7 days after CP treatment) TOV513 (top) and TOV1294G (bottom) cells during treatment with 5 µM of ABT-263 for 72 hours. Statistical significance was calculated using the unpaired t test. * p < 0.05; ** p < 0.01, *** p < 0.001, **** p < 0.0001.

### HGSOC tissues display senescence hallmarks following exposure to chemotherapy in patients

Given that TIS occurs in HGSOC primary cultures in response to chemotherapy regimens used in the clinic, we explored whether TIS also occurs in patients as a response to treatment. We constructed a tissue microarray (TMA) from HGSOC tissues collected from 170 patients treated at the CHUM: 85 patients who had undergone surgery prior to chemotherapy, providing chemotherapy naïve samples (pre-chemotherapy), and 85 patients who had undergone surgery after primary chemotherapy, providing post-chemotherapy samples. Clinical follow-up data was associated with each patient sample (Fig. S5). The TMA was stained using IF to detect SA biomarkers, digitalized at high resolution, and quantitatively analyzed. Tissue cores were identified, segmented into epithelial (i.e., cancer cells) and stromal compartments using epithelial markers (cytokeratin CK7, 18, and 19), further segmented into nuclear and extra-nuclear compartments using DAPI (nuclear DNA signal), and categorized in specific compartments (i.e., total epithelial, epithelial nuclear, total stromal, and stromal nuclear; Fig. 5a). As expected for ovarian cancer (50), chemotherapy caused a significant reduction in the relative epithelial surface area within tumor tissues (Fig. 5b) also reflected by the overall diminishment in total core mean fluorescent intensity (MFI) for cytokeratins or E-Cadherin (epithelial), and increased vimentin (stromal) (Fig. 5c-e). We first explored senescence biomarkers in tissue compartments using matched pre- and post-chemotherapy samples collected from the same patients, termed paired samples; these were of limited availability. Interestingly, the MFI of epithelial nuclear lamin B1 and Ki67 consistently decreased, while total epithelial IL6 consistently increased post-chemotherapy (Fig. S6a-c, g-i), consistent with changes expected in the context of HGSOC senescence based on primary culture results. Similar trends were seen in the stromal compartment as well, albeit less consistently (Fig. S6a-c, g-i). Trends in IL8, vimentin, E-Cadherin,p16^INK4A^, PML, MCM2 and Geminin were less clear in both the epithelium and the stroma (Fig S6d-f, j-m; n, s). To gain a broader view, we examined SA biomarkers MFI variations in the cohort as a whole. Consistent with a TIS response to treatment in HGSOC epithelium, epithelial nuclear Ki67, geminin and lamin B1 decreased significantly in post-chemotherapy samples, whereas epithelial nuclear PML and total epithelial IL6 increased (Fig. 5f). There was no change in epithelial apoptosis levels as evaluated using caspase-3 cleavage as a measure of activation (Fig. 5f, S6o, t for paired samples). Interestingly, total epithelial vimentin and E-Cadherin were found to increase post-chemotherapy (Fig. 5f), suggesting an epithelial-to-mesenchymal transition (51), a phenomenon also linked to paracrine effects of the SASP (8, 52). In contrast, and unlike results observed in primary cultures, no significant differences were observed for epithelial nuclear p16^INK4A^ and MCM2 or total epithelial IL8 between pre- and post-chemotherapy samples (Fig. 5f). Similarly, trends in the MFI of several SA biomarkers were consistent with senescence in the tumor stroma. In particular, stromal nuclear Ki67, geminin and lamin B1 decreased, while stromal nuclear PML and total stromal IL8 increased (Fig. 5g). Unlike epithelial tumoral tissue, we observed slightly increased apoptosis rates in the stroma, but no significant difference was found in stromal p16^INK4A^, MCM2, IL6, or vimentin, between pre- and post-chemotherapy tissues (Fig. 5g).

**Figure 5:**
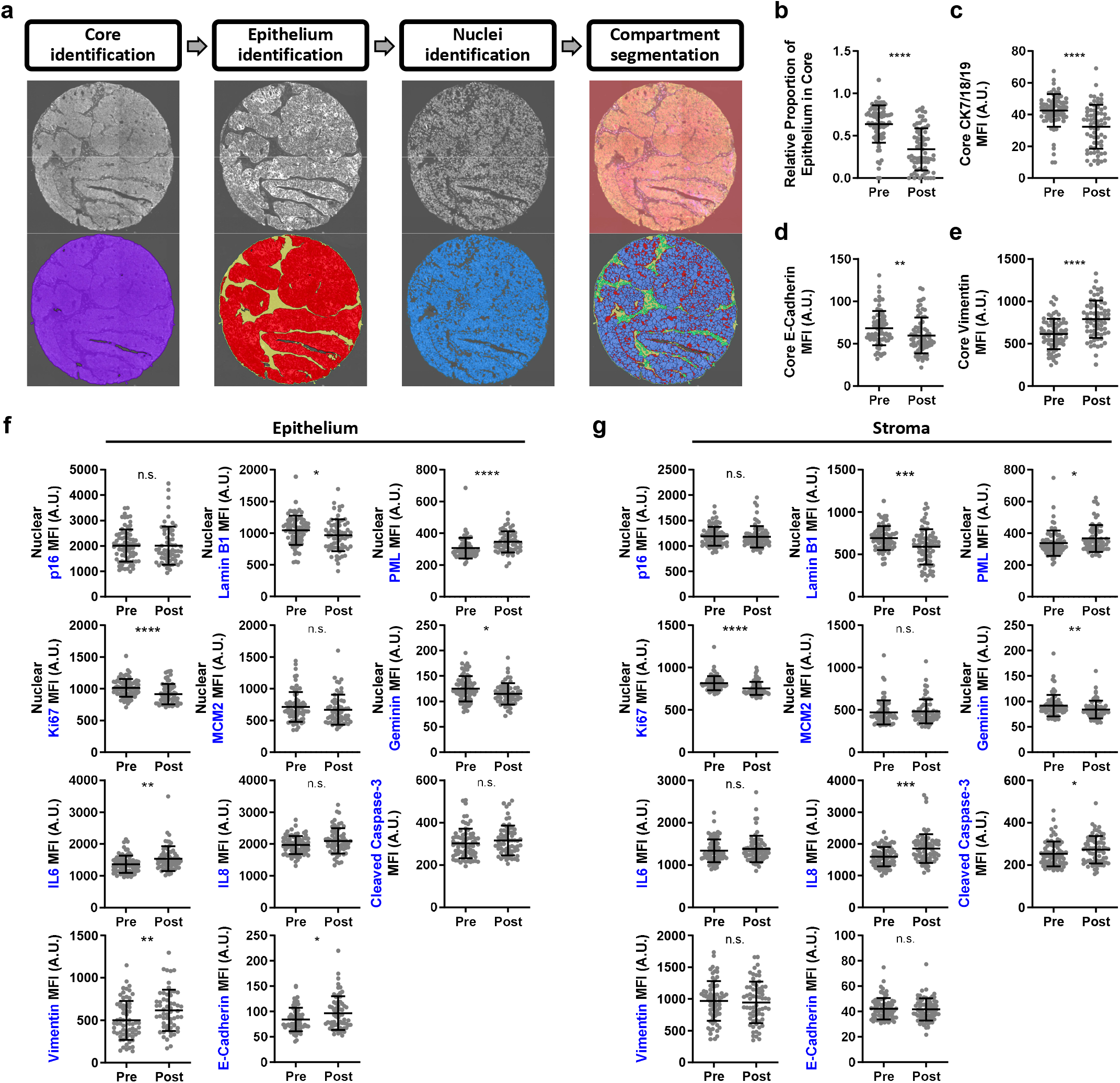
HGSOC tissues display senescence hallmarks following exposure to chemotherapy in patients. (**a**) Segmentation protocol of multi-channel IF images from a representative HGSOC tissue core. (**b-e**) Quantification of the proportion of epithelium in pre- and post-chemotherapy cores via the ratio of epithelium to core (**b**), the total core MFI of the epithelial mask (cytokeratins 7, 18, 19) (**c**), the total core E-cadherin MFI (**d**), and the total core vimentin MFI (**e**). (**f**) Quantification of the MFI of senescence markers (from left to right, top to bottom: nuclear p16, nuclear lamin B1, nuclear PML, nuclear Ki67, nuclear MCM2, nuclear geminin, IL6, IL8, cleaved caspase-3, vimentin, and E-cadherin) in the epithelium of pre- and post-chemotherapy tissue cores. (**g**) Quantification of the MFI of senescence markers (from left to right, top to bottom: nuclear p16, nuclear lamin B1, nuclear PML, nuclear Ki67, nuclear MCM2, nuclear geminin, IL6, IL8, cleaved caspase-3, vimentin, and E-cadherin) in the stroma of pre- and post-chemotherapy tissue cores. Statistical significance was calculated using the Mann Whitney test. n.s., not significant; * p < 0.05; ** p < 0.01, *** p < 0.001, **** p < 0.0001.

### Single cell HGSOC tissues analysis reveals senescence-associated cell cycle perturbations

In order to deepen our understanding of cellular responses in HGSOC tissues, we next sought to determine the distribution of single cells in the cell cycle in each core, paying particular attention to the proportion of cells having exited the cell cycle. To this end, we analyzed co-stainings of cell proliferation markers MCM2 and Ki67, as well as MCM2 and geminin, which are differentially expressed during the cell cycle (Fig. 6a). MCM2 is highly expressed in G_1_ and less strongly expressed in S, G_2_ and M (Maiorano et al. 2006). On the other hand, Ki67 is weakly expressed in G_1_ and early S phase, and more strongly expressed in later phases of the cell cycle (Li et al. 2014). Similarly, geminin is expressed as of the G_1_-S transition and its expression increases throughout the rest of the cell cycle (Wohlschlegel et al. 2002; Loddo et al. 2009). Importantly, none of the three markers are expressed in G_0_ (Gerdes et al. 1984; Maiorano et al. 2006; Loddo et al. 2009). Individual nuclei were identified in the epithelial and stromal compartments and classified according to MCM2 and Ki67 or geminin expression (Fig. 6b). In line with the occurrence of TIS in response to chemotherapy, post-treatment samples displayed a lower cell density in the epithelial compartment, suggesting an increase in cell size in this compartment (Fig. 6c). Interestingly, cell density was increased in the stromal compartment (Fig. 6c). Importantly, however, cell cycle classification of cells in the epithelial compartment revealed a significant increase in the proportion of cells having exited the cell cycle in post-chemotherapy tissues in both the Ki67-MCM2 and Geminin-MCM2 analyses, lending further support for the occurrence of TIS in response to chemotherapy (Fig. 6d). Conversely, the proportion of cells in G_1_, S, and G_2_/M showed either a trend or a significant decrease, with a particularly notable decrease in the proportion of cells positive for MCM2 and Ki67 or geminin (i.e. cells in S phase) (Fig. 6d). Similar, though generally less ample variation were observed in the stromal compartment (Fig. 6d). Overall, these findings support the notion that TIS occurs as a response to treatment in HGSOC cells *in vitro* and in patient, suggesting TIS is an important response to current HGSOC chemotherapy treatment.

**Figure 6:**
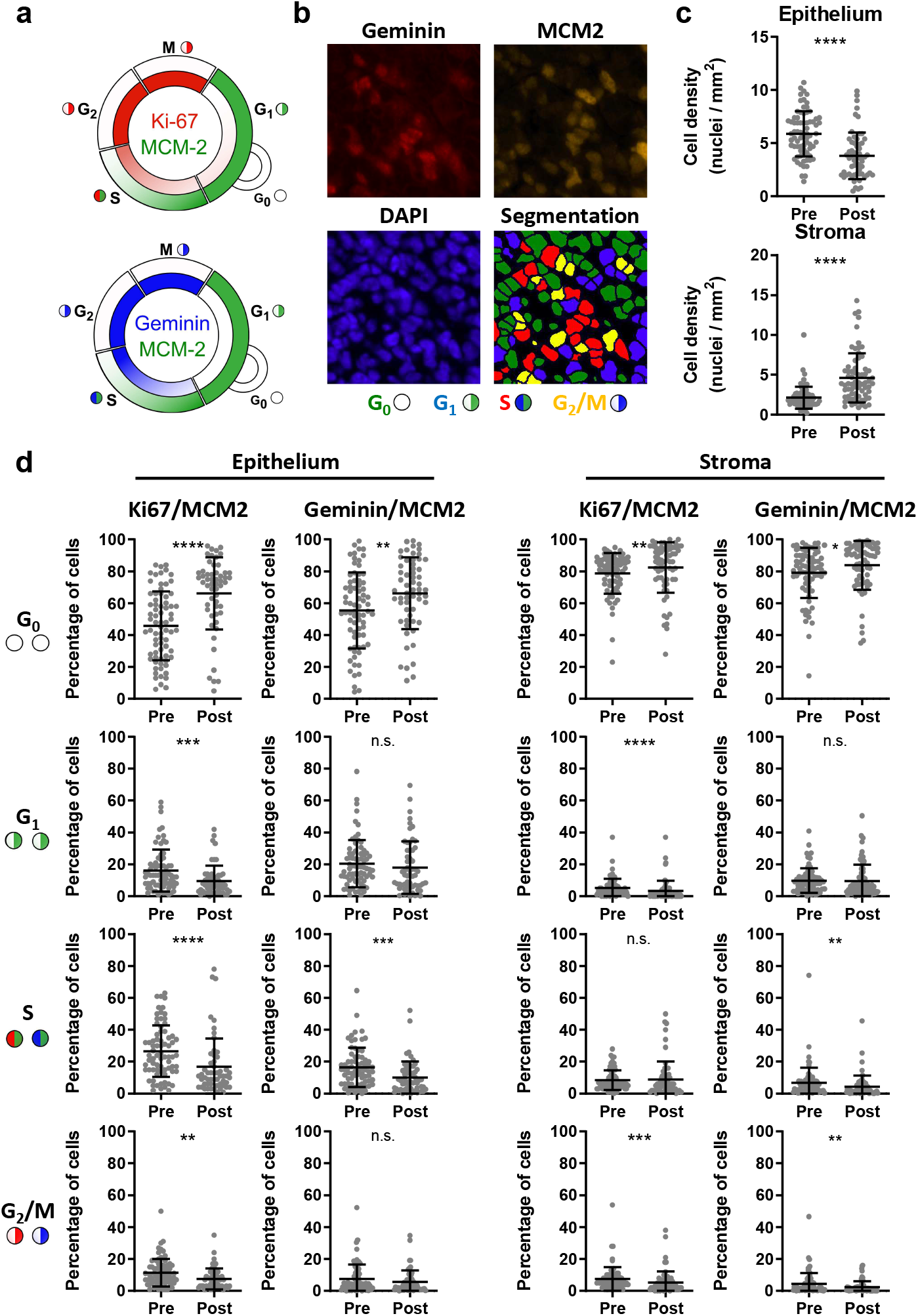
Cells in HGSOC tissues exit the cell cycle following exposure to chemotherapy in patients. (**a**) Expression patterns of MCM2, Ki67 and geminin in G0, G1, S and G2/M. See text for details. (**b**) Classification of individual nuclei in different phases of the cell cycle from a representative HGSOC tissue core according to MCM2 and geminin expression (G0: MCM2- /geminin-; G1: MCM2+/geminin-; S: MCM2+/geminin+; G2/M: MCM2-/geminin+). (**c**) Density of nuclei expressed as nuclei per mm2 in pre- and post-chemotherapy tissue cores, in the epithelial (top) and stromal (bottom) compartments. (**d**) Proportion of cells in G0, G1, S and G2/M (ordered from top to bottom) according to MCM2 and Ki67 or geminin expression in the epithelial (left) and stromal (right) compartments in pre- and post-chemotherapy HGSOC tissue cores. Statistical significance was calculated using the Mann Whitney test. n.s., not significant; * p < 0.05; ** p < 0.01; *** p < 0.001; **** p < 0.0001.

### Senescence-associated biomarker levels in post-chemotherapy HGSOC tissues correlate with 5-year survival

Whether senescence is beneficial or detrimental during human cancer treatment remains unknown. To determine how senescence and TIS may influence patient’s clinical course, we split all the patients from our clinical cohort regardless of chemotherapy status, as well as patients specifically from the pre- and post-chemotherapy groups, into high and low expression of SA biomarkers and compared 5-year overall survival between the two groups using Kaplan-Meier analysis (Fig. S7a-b). Cutoff values for high- and low-expression were determined via receiver operating characteristics (ROC) analyses. Low expression of lamin B1 and high expression of p16^INK4A^ were each associated with improved 5-year overall survival in the post-chemotherapy subgroup, suggesting the occurrence of TIS post-chemotherapy to be beneficial to clinical outcomes (Fig. 7a-b). Interestingly, this association was maintained for p16^INK4A^ in the pre-chemotherapy subgroup, as well as for both markers in our entire cohort, independently of chemotherapy status, suggesting senescence at large may be beneficial to clinical outcomes in HGSOC (Fig. 7a-b). Importantly, stratification of patients according to both p16^INK4A^ and lamin B1 (i.e. high p16^INK4A^ and low lamin B1 vs. low p16^INK4A^ and high lamin B1), which likely more specifically identified tissues with senescence, strengthened the associated between ameliorated 5-year overall survival and biomarkers suggestive of TIS or senescence in the whole cohort and in the post-chemotherapy group specifically (Fig. 7c). Stratification of patients according to the distribution of cells in the cell cycle according to MCM2 and geminin status yielded similar results. A high proportion of epithelial cells in G_0_ was associated with a non-significant trend towards increased 5-year overall survival in the whole cohort and in the post-chemotherapy group (Fig. 7d). Conversely, a low proportion of epithelial cells in G_2_/M was associated with improved 5-year overall survival across all groups analyzed (Fig. 7e). Finally, stratification of patients according to proportion of cells in either G_0_-G_1_ or S-G_2_/M showed a trend towards improved outcomes with a high proportion of cells in G_0_-G_1_ and a low proportion of cells in S-G_2_/M in both the pre-and post-chemotherapy groups, and a significant association in the whole cohort (Fig. 7f). Analyses in the stromal compartment did not yield consistent trends in any of the subgroups or in the whole cohort (Fig. S7b). Overall, our data points to a clinical benefit to TIS occurring post-chemotherapy in the epithelial compartment, as well as a clinical benefit to senescence at large in HGSOC tissues.

**Figure 7:**
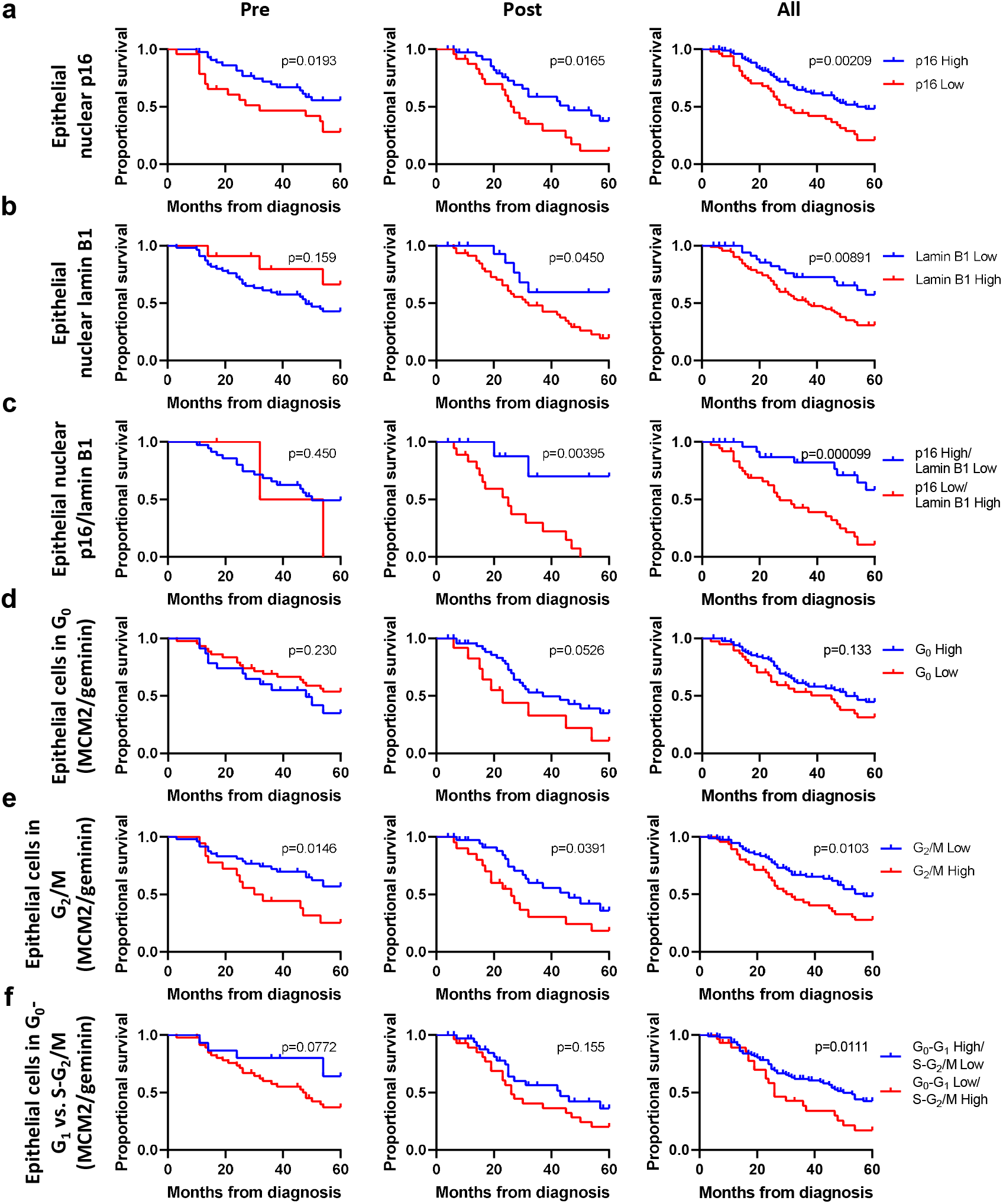
Senescence-associated biomarker levels in post-chemotherapy HGSOC tissues correlate with 5-year survival. (**a**) Kaplan-Meier analysis comparing 5-year survival between high and low expression of p16INK4A before (n=68) and after (n=63) chemotherapy, as well as independently of chemotherapy status (n=131) (**b**) Kaplan-Meier analysis comparing 5-year survival between high and low expression of lamin B1 before (n=67) and after (n=65) chemotherapy, as well as independently of chemotherapy status (n=132). (**c**) Kaplan-Meier analysis comparing 5-year survival between high expression of p16INK4A with low expression of lamin B1 and low expression of p16INK4A with high expression of lamin B1 before (n=39) and after (n=29) chemotherapy, as well as independently of chemotherapy status (n=65). (**d**) Kaplan-Meier analysis comparing 5-year survival between high and low proportion of cells in G_0_ (according to MCM2 and geminin expression) before (n=67) and after (n=59) chemotherapy, as well as independently of chemotherapy status (n=126). (**e**) Kaplan-Meier analysis comparing 5-year survival between high and low proportion of cells in G_2_/M (according to MCM2 and geminin expression) before (n=67) and after (n=59) chemotherapy, as well as independently of chemotherapy status (n=126). (**f**) Kaplan-Meier analysis comparing 5-year survival between high and low proportion of cells in G_0_-G_1_ (i.e. respectively low and high proportion of cells in S-G_2_/M) (according to MCM2 and geminin expression) before (n=62) and after (n=65) chemotherapy, as well as independently of chemotherapy status (n=127). Groups were separated into high and low espressors or proportions based on cut offs determined by receiver operating characteristics (ROC) analyses. Lamin B1 and p16INK4A expression status determined separately for each marker was used to determine p16INK4A high/lamin B1 low and p16INK4A low/lamin B1 high groups. Statistical significance was calculated using the Log Rank test.

## DISCUSSION

In this study, we demonstrate that HGSOC primary cells taken from highly aggressive and fatal human cancers respond to chemotherapy by triggering TIS, including most senescence hallmarks. Senescence is often driven by the p53/p21^WAF1^ and p16^INK4A^/Rb signaling cascades, but TP53 is rendered dysfunctional through a multitude of mechanisms in nearly all HGSOC (24, 39, 40, 43). It is thus perhaps not surprising that we found chemotherapy (CP) and IR promote p16^INK4A^-associated HGSOC senescence. Indeed, the key senescence tumor suppressor p16^INK4A^ is rarely mutated in HGSOC (Fig. S2c), and its diffuse expression is used by pathologists to help differentiate HGSOC from other ovarian cancer subtypes (53). Our results raise the intriguing possibility that HGSOC cells often maintain at least some p16^INK4A^ functions for cell cycle control in the absence of other TP53-associated DNA repair checkpoints, simultaneously preserving a capacity to undergo senescence in response to acute genome stress. A parallel non-exclusive possibility is that the p16^INK4A^/Rb pathway is inherently subverted downstream of p16^INK4A^ in HGSOC, as often happens in HPV-induced cancers (54), which would also explain low mutation rates of *CDKN2A*. Supporting both possibilities, our attempts to immortalize HGSOC cells through simple shRNA knockdown of p16^INK4A^ failed in both primary cultures we tested (data not shown), suggesting that robust SAPA pathways other than the traditional p16^INK4A^/Rb and p53/p21^WAF1^ cascades remain to be identified in this context. Nevertheless, the evidence presented here of p16^INK4A^-associated replicative senescence (in most of our primary cultures) combined with the low *CDKN2A* mutation prevalence in HGSOC patients (3-5%) suggests that the majority of HGSOC tumors retain the ability to undergo TIS, providing treatment enhancement opportunities based on pharmacological senescence manipulation.

In general, ethical and clinical considerations preclude the collection of solid tumor tissue during chemotherapy regimens. HGSOC presents a particular clinical opportunity in that patients undergo either neoadjuvant chemotherapy followed by interval debulking surgery or primary debulking surgery followed by adjuvant chemotherapy (55). These two treatment trajectories can provide access to either pre-or post-chemotherapy samples. Our comparison of tissues from patient cohorts that have undergone either treatment trajectory suggests that TIS occurs in ovarian cancer tissues in response to chemotherapy, and the associated clinical follow-up data further reveal that TIS may be beneficial in this context. This is important, as the activation of TIS and SASP following chemotherapy in HGSOC primary cultures and patient tissues suggest that ovarian cancer DNA-damaging chemotherapy can alter the post-treatment tumoral tissue microenvironments, further guiding potential pharmacological avenues toward the targeting of senescence via SAPA reinforcement, SASP manipulation, and senescent cancer cell elimination by senolysis (7, 10, 20).

At the same time, several observations were made that raise unanswered questions. In particular, why is p16^INK4A^ upregulation readily observed in all TIS primary HGSOC cultures but not observed in TIS patient tissues? It is important to note that post-chemotherapy tissues were collected a minimum of four weeks after the most recent chemotherapy cycle, as per standard clinical protocols, to allow for patient recovery before surgery. During this period, it is possible that cells that upregulate p16^INK4A^ relative to the baseline expression had indeed become senescent, but may have been subjected to immune clearance (21, 56), reverted from their senescence-like state as we have shown for HGSOC cell lines (26), or may have been lost by dilution to cell populations that have not efficiently activated TIS. In this context, it is also important to note that the post-treatment samples available for study are necessarily taken from patients with incomplete responses to chemotherapy, i.e. with resectable tumour after chemotherapy. Though only a minority of patients undergoing neoadjuvant chemotherapy will have no resectable tumour at the time of interval debulking surgery, this proportion is not insignificant and is several fold greater than among those undergoing primary debulking surgery (55). In accordance with this selection of patients with poorer responses to chemotherapy, overall survival is inferior in the post-chemotherapy group (i.e. patients with resectable tumour at interval debulking surgery) as compared to the pre-chemotherapy group (supplementary figure 8). Thus, it remains possible that tissues that have undergone a strong senescence response and subsequent immune clearance were underrepresented in this study. Conversely, it is probable that tissues underwent cell fates other than senescence as well, such as apoptosis, although our data suggest dying cells would also have already disappeared when we probed the tissues given that apoptosis levels were relatively unchanged pre/post-treatment. Taken together, a closer study of the immediate post-treatment period would be crucial in order to improve our understanding of HGSOC TIS, but it will require innovative clinical strategies to obtain the required human tissues.

Overall our data suggests that current chemotherapy regimens are efficient at inducing TIS in senescence competent HGSOC cells, but that this TIS induction is probably suboptimal. It is likely that a subset of cancer cells either avoid or escape the senescence cell fate to potentially emerge following treatment. In light of this, the enhancement or reactivation of senescence in these cells may be an interesting and novel strategy in order to harness the curative potential of TIS in HGSOC. Perhaps as important, the presence of biologically versatile senescent cells in a treated tumor implies the possibility that these influence other cells within the tumor in ways that alter treatment outcomes. For example, the DNA damage-induced secretion of IL6 in the senescent microenvironment of post-chemotherapy murine thymus is sufficient to create a protective niche for a subset of lymphoma cells that can cause cancer recurrence (22). Similarly, the presence of senescent cells in post-therapy murine breast cancers has been shown to trigger increased SASP levels which are associated with cancer recurrence (57). Finally, the secretion of the SASP factor WNT16B (wingless-type MMTV integration site family member 16B) by senescent stromal cells in post-therapy murine prostates provides direct support for continued growth of prostatic epithelial cancer cells. Notably, increased stromal WNT16B levels have also been detected in post-therapy human prostate, ovarian, and breast cancers (52). Provided we report alternate evidence that senescence could be beneficial during human HGSOC treatment, it is clear that senescence and its relationship with long-term treatment outcomes must be assessed in a context-dependent manner associated with key clinical data. Except for the emerging vision presented here for HGSOC, this concept has yet to be thoroughly studied in human cancers.

In the case of HGSOC, given the unpredictable influence of senescent cells on the clinical course, our previous report in HGSOC cell lines combined to the results presented here in a context closer to HGSOC patient support the idea that the incorporation of senolytic drugs into treatment regimens could prove to be beneficial via enhanced elimination of senescent cells. Along the same lines, prior to senolytic clearance, using small molecule CDKi or pharmacologic epigenetic regulators could be considered to strengthen p16^INK4A^-induced suppression of the cell cycle as previously explored for ovarian and prostate cancer (58, 59). Overall, results from this study support the notion that senescence is a therapeutic target to develop novel treatment modalities in HGSOC.

## MATERIALS AND METHODS

### Patients and tissue specimens

Tumor samples were obtained from patients who underwent surgery at the CHUM Department of Gynecologic Oncology for ovarian cancer between 1993 and 2014. Disease stage was defined using the International Federation of Gynecology and Obstetrics (FIGO) staging system, and tumor histopathology was classified using World Health Organization (WHO) criteria. Patient survival was calculated from the date of diagnosis, which corresponded to 3 months prior to surgery for post-chemotherapy samples. Five-year overall patient survival was defined as the time from diagnosis to death or last follow-up, up to a maximum follow-up of 5 years.

### Pre- and post-chemotherapy epithelial ovarian tumor tissue microarray (TMA)

A gynecologic pathologist reviewed all cases. The type of ovarian carcinoma was identified and areas of interest were marked on hematoxylin and eosin–stained slides. Two cores of 0.8-mm diameter for each tissue sample were arrayed onto recipient paraffin blocks. Although the final TMA was composed of 340 cores from 170 epithelial ovarian tumors (2 replicate cores per patient), the usable cores per analysis is detailed in sTable2. Clinicopathological details of the cohort are summarized in Fig. S4.

### Antibodies

Details of primary and secondary antibodies used in this study are summarized in sTable3.

### Immunofluorescence (IF) staining of TMA

The TMA was sectioned into 4-μm thick slices and processed/stained using a BenchMark XT automated stainer (Ventana Medical System Inc., Tucson, AZ, USA). Antigen retrieval was carried out with Cell Conditioning 1 (Ventana Medical System Inc.; no. 950-124) for 60 minutes. The prediluted primary antibody was automatically dispensed and incubated for 60 minutes at 37°C. The following steps were performed manually. After blocking with Protein Block serum-free reagent (Dako #X0909), secondary antibodies were added for 45 minutes, followed by washing and blocking overnight at 4°C with diluted Mouse-On-Mouse reagent (Vector #MKB-2213). This was followed by an incubation of 60 minutes with primary antibodies of the epithelial mask. The Alexa Fluor 750 secondary antibody was incubated for 45 minutes, followed by DAPI staining for 5 minutes. Staining with 0.1% Sudan black for 15 minutes to quench tissue autofluorescence was followed by coverslip mounting using Fluoromount Aqueous Mounting Medium (#F4680, Sigma). The full TMA was scanned-digitalized using a 20X 0.75NA objective with a resolution of 0.325 µm (BX61VS, Olympus). Multicolor images were segmented and quantified using Visiopharm Integrator System (VIS) version 4.6.1.630 (Visiopharm, Denmark).

### VIS analysis

IF scoring was performed using the VIS software. Briefly, VIS was used to measure the MFI of all pixels in original digitalized multicolor TMA images for a chosen region of a tissue sample (total tissue core or core sub-compartments as described for Fig. 5a). Then, the MFIs corresponding to replicate total cores or selected core sub-compartments from the same patient are averaged to calculate one final mean MFI for each data point used in the presented analysis. Controls related to duplicate core reproducibility are presented in sFig. 9. VIS was used to perform analysis of the cell cycle distribution of cells in HGSOC tissues. Individual nuclei in each core were identified and assigned phase of the cell cycle (G_0_, G_1_, S, or G_2_/M) according to the MFI of MCM2 and either Ki67 or geminin for that particular nucleus. Proportion of nuclei in each phase of the cell cycle in replicate cores from the same patient were averaged to calculate a final proportion for each data point used in the present analysis.

### Statistical analysis for TMA

Statistical significance of differences in MFI between pre- and post-chemotherapy groups were calculated using the Mann Whitney test. Survival curves were plotted using Kaplan-Meier analyses and the log-rank test was used to test for significance. All statistical analyses were done using the Statistical Package for Social Sciences software version 21 (SPSS, Inc.), and statistical significance was set at p<0.05. Some patients were excluded from final analysis based on poor quality cores (folded or scratched cores) or for lack of epithelial material in cores (the usable cores per analysis are detailed in sTable2).

### Primary cell culture

Associated clinical data were available for all HGSOC primary cells (TOV) and primary normal epithelial ovarian cells (NOV) used. All primary cancer cells used were HGSOC with FIGO stage II-IV. Primary cell cultures were derived from patient samples following procedures reported previously (60). Unless otherwise indicated, cells were cultured in OSE media supplemented with 15% fetal bovine serum (FBS) and 1% streptomycin, in a 5% O_2_ incubator.

### SA-β-gal staining

Senescence-associated β-galactosidase (SA-β-gal) was assessed as previously described (61). Pictures were taken with a Nikon Eclipse TE300 microscope (Nikon Instruments Inc. Melville, NY, U.S.A).

### Irradiation and chemotherapy

Cells were seeded in appropriate vessels and 48 hours after seeding, cells were treated with either 10 Gy of ionizing radiation (IR) using a Cesium 137 irradiator, or a combination of carboplatin (10 μM) and paclitaxel (30 nM). To mimic dosage used in HGSOC patients Carboplatin and paclitaxel were removed after 12 hours (44, 45). Senescence hallmarks were tested 8-10 days later as indicated for individual experiments.

### Immunofluorescence staining of primary cell cultures

Cells were cultured in 4- or 8-well chamber slides (Nunc, Penfield, NY, USA). After treatment and incubation, cells were fixed in 10% formalin (Sigma) for 10 minutes and washed 3 times in PBS for 5 minutes at room temperature (RT), followed by incubation for 30 minutes in blocking buffer (1% IgG-free BSA, 4% donkey serum (Jackson ImmunoResearch, West Grove, PA, USA) in PBS), then by incubation with primary antibodies diluted in blocking buffer overnight at 4 °C. Cells were then washed and incubated with secondary antibodies for 1 hour at RT. Finally, slides were washed and mounted with ProLong Gold Antifade reagent with DAPI (Life Technologies, Willow Creek Road Eugene, OR, USA). Images were acquired using a Carl Zeiss Axio Observer Z1 fluorescence microscope with AxioVision software (45 Valley brook Drive Toronto, ON, Canada) and presented with Photoshop CS (Adobe, San Jose, CA, USA). Images were then either analyzed visually or via computer-assisted quantitative image analysis. Computer-assisted quantitative image analysis was performed on raw images using the AxioVision ASSAYbuilder software (45 Valley brook Drive Toronto, ON, Canada), Physiology Analyst module. Cell nuclei were identified and delimited using the DAPI signal. For EdU and p16, the EdU and p16 signal MFI in each cell nucleus was quantified. Cut-offs for positive and negative EdU and p16 staining were determined subjectively based on the lowest value that deviated from background noise in each cell culture. For lamin B1, the MFI of the lamin B1 signal was quantified in a 6 µM-wide band along the periphery of the nucleus, stretching from the outer limit of the nucleus inwards. Unless otherwise specified in the figure legends, each data point represents the average value measured in one cell culture. Computer-assisted image analysis was used for cultures TOV1294G, TOV2267 and TOV2356 for p16, EdU and lamin B1 quantification. Quantification of the proportion of p16-positive and EdU-postitive cells in cultures TOV513 and TOV959D was performed visually. Quantification of the proportion of cells with 3 or more 53BP1 foci was performed visually for all primary cell cultures.

### DNA synthesis detection

Cells in chamber slides were pulsed with the modified thymidine analogue EdU for 48 hours before fixation. Cells were processed as per the manufacturer’s protocol to detect the incorporation of EdU into DNA using Click-iT chemistry (Invitrogen C10340) and Alexa Fluor 488 azide (No. A10266) or Alexa Fluor 647 azide (No. A10277). Following washing with PBS, IF staining for p16INK4A was performed as described above. Finally, cells were washed with PBS, mounted with ProLong Gold Antifade reagent with DAPI (Life Technologies, Willow Creek Road Eugene, OR, USA), and analyzed as for IF above.

### Gene expression analysis

RNA from HGSOC cultures were extracted and purified as previously described (62) and microarray experiments were performed at the McGill University and Genome Quebec Innovation Centre (genomequebec.mcgill.ca) using HG-U133A GeneChip arrays (Affymetrix®, Santa Clara, CA). Gene expression levels were calculated for each probe set from the scanned image by Affymetrix GeneChip (MAS5) as previously described (62). As reproducibility of expression data is variable at low raw expression values(63), normalized data points below 15 were reassigned a threshold value of 15 based on the mean expression value of the lowest reliability scores. Probe sets having these threshold values in all samples were not used for further analysis. Normalized gene expression data from 42 HGSOC primary cultures (sTable 1) was used to stratify expression clusters using unsupervised k-means (Gene Cluster 3.0) based on 177 E2F-target genes (Genes encoding cell cycle-related targets of E2F transcription factors, M5925, http://software.broadinstitute.org/gsea/msigdb/cards/HALLMARK_E2F_TARGETS). Clustering results were generated using Java Treeview to highlight E2F-response clusters I, II, III; cluster I reflects high proliferation (high E2F-activity) and cluster II and III reflects lower EF2-activiy (Fig. s1a-b). Early passage primary cultures from cluster I (≤50% replicative lifespan completed; n=5) were then compared to late passage primary cultures from cluster II-III (≥75% replicative lifespan completed; n=7) and the differential gene expression detected between these 2 groups (early versus late passage) were compared using GSEA to 22 published senescent signatures (64) (Fig. s1c-d).

### SASP profiling

SASP was prepared as previously described (65) with the following adjustments: cells were seeded and treated with paclitaxel/carboplatin as described above, followed by incubation in OSE medium with 15% FBS for another 8 days (media was changed every 2 days). Sixteen hours before performing the antibody array, the vessels were washed 3 times with PBS, and the media was replaced with serum-free media to generate serum-free conditioned media. Fresh conditioned media was collected and stored on ice for immediate use on multiplex ELISA, and cell numbers were counted. Multiplex secretome detection was performed according to the protocol provided by Meso Scale Discovery V-PLEX Human Biomarker (40-Plex Kit K15209D-2). In brief, 50-μl samples, diluted 1:2 or 1:4, were added to the wells. Plates were sealed and incubated for 2 hours at RT. After incubation, plates were washed 3 times with 150 μl of wash buffer per well, and 25 μl of detection antibody solution were then added to each well. The plates were incubated for another 2 hours at RT, and washed 3 times. Before reading, 150 μl of 2X read buffer T were added to each well. Finally, the plates were read on the MSD instrument, with signals normalized to cell number.

### Measurements of relative growth and viability

Cells were seeded in 96-well plates and cultured within an IncuCyte Zoom live cell imager that captured phase contrast images every 2 hours (Essen Bioscience). Twenty-four hours after seeding, cells received genotoxic treatments as indicated. Using captured images, the relative cell confluence (confluence mask) was determined by an integrated confluence algorithm (Essen Bioscience) as previously done (26). To track treatment response, the data is reported as relative cell confluence (fold change) representing the confluence during or at the end of the indicated treatments over confluence just before treatment is initiated.

### Study approval and ethics statement

Approval from the CHUM institutional ethics committee was obtained. Informed consent from all patients was obtained before sample collection and samples are anonymized.

## ACKNOWLEDGEMENTS

We thank members of the Mes-Masson, Provencher and Rodier laboratory for valuable comments and discussions, as well as Suzana Anjos and Jacqueline Chung for language editing. We thank the CRCHUM molecular pathology core facility and the Institut du cancer de Montréal (ICM) Imaging and Live imaging platform. This work was supported by the ICM (DP, AMMM, FR) and by the Canadian Institute for Health Research (CIHR MOP114962 to FR), the Terry Fox Research Institute (TFRI 1030 to FR) and the Cancer Research Society (CRS) in partnership with Ovarian Cancer Canada (20087 to AMMM, DP; 22713 to FR). Microarray data was generated using funding from Genome Canada/Genome Québec (to AMMM). AMMM, DP and FR are researchers of CRCHUM/ICM, which receive support from the Fonds de recherche du Québec - Santé (FRQS). FR is supported by a FRQS junior I-II career awards (22624, 33070). Ovarian tumor banking was supported by the Banque de tissus et de données of the Réseau de recherche sur le cancer of the FRQS affiliated with the Canadian Tumor Repository Network (CTRNet). LCG and MS received Canderel fellowships from the ICM, SC received a FRQS postdoctoral award, MS received a CIHR doctoral award and YZ was supported by an ICM/MITACS postdoctoral fellowship.

## AUTHORS CONTRIBUTIONS

LCG, SC, MS and FR designed the study. LCG, SC, IC, MS, and JL performed experiments. YZ and EC performed bioinformatics analysis, KR revised all pathology samples for TMA construction, LCG, SC, MS, IC, and FR collected and analyzed data. LP and MD performed tissue banking and provided technical assistance. LCG, SC, MS, and FR wrote the manuscript. AMMM and DP provided technical support, biobank access, molecular pathology expertise, conceptual advice, and revised the manuscript.

---

**Supplemental figure 1: Proliferative lifespan transcriptome analysis of primary HGSOC cultures.** (**a**) Major steps in the workflow of the analysis of transcriptome data from HGSOC primary cell cultures. Transcriptome data was collected via Affymetrix microarrays from 42 HGSOC primary cell cultures, each at a different passage with respect to total proliferative lifespan of individual cultures. Unsupervised clustering was then performed according to E2F target gene expression level, yielding 3 clusters shown in (**b**). Using the eventual maximal passage number for each primary cell culture, cultures were selected from clusters 1 and 3, respectively, if the passage at which transcriptome data was acquired was 50 % or less and 75 % or more of the maximal eventual passage number of that culture as shown in (**c**). These cell cultures were considered to be “early passage” and “late passage” respectively. Late passage cultures were compared to early passage cultures on gene set enrichment analysis (GSEA) using gene sets generated by Hernandez-Segura et al. (2017) and downloaded from https://www.ncbi.nlm.nih.gov/geo/. (**b**) Unsupervised clustering of 42 HGSOC primary cell cultures according to E2F target gene expression. (**c**) Primary cell cultures included in the “early passage” (n=5) and “late passage” (n=7) groups. Primary cell cultures were selected from clusters I (high proliferation) and II-III (low proliferation) in (**b**), respectively, if the passage at which transcriptome data was acquired was 50 % or less and 75 % or more of the maximal eventual passage number of that culture, in order to form the early and late passage groups. (**d**) Different gene sets described by Hernandez-Segura et al. (2017) significantly upregulated in “late passage” (left) and “early passage” (right) cell cultures on GSEA.

**Supplemental figure 2: Normal and HGSOC primary cultures undergo proliferative senescence.** (**a**) Early- and late-passage normal ovarian epithelial cell (NOV1341G) primary cultures were stained for SA-β-Gal activity. (**b**) Early- and late-passage HGSOC (TOV959D, TOV1294G, TOV2267, and TOV2356) primary cultures were labelled with EdU (24-hour pulse) (representative TOV2356 images shown). (**c**) Early- and late-passage normal ovarian epithelial cell (NOV1341G) primary cultures were labelled with EdU (24-hour pulse). (**d**) Mutation rates for *TP53* and *CDKN2A* found in two large The Cancer Genome Atlas (TCGA) ovarian cystadenocarcinoma cohorts, including 311 cases (provisional) and 316 cases (Cancer Genome Atlas Research, 2011), using cBioportal. (**e**) Quantification of the relative mean fluorescent intensity (MFI) of nuclear lamin B1 per cell from (Fig. 1 h) (each cell represents a data point); statistical significance was calculated using the unpaired t test. **** p < 0.0001.

**Supplemental figure 3: HGSOC primary cultures undergo TIS in response to DNA damage and chemotherapy.** (**a**) Quantification of cells with more than three 53BP1 foci in TOV513, TOV1294G, TOV2267, and TOV2356 cultures in either untreated conditions or after IR or CP treatment (each high-powered field represents a data point). (**b**) Quantification of lamin B1 MFI per cell normalized to control in TOV1294G, TOV2267, and TOV2356 after IR or CP treatment (each cell represents a data point). Statistical significance was calculated using the unpaired t test. ** p < 0.01; *** p < 0.001, **** p < 0.0001.

**Supplemental figure 4: HGSOC TIS renders cells sensitive to ABT-263.** (**a-b**) Quantification of the relative confluence of non senescent TOV513 (a) and TOV1294G (b) cells after treatment with various concentrations of ABT-263 for 72 hours. Statistical significance was calculated using the unpaired t test. * p < 0.05; ** p < 0.01, *** p < 0.001, **** p < 0.0001.

**Supplemental figure 5: Clinical characteristics of the patients in the pre- and post-chemotherapy HGSOC cohorts.** (**a**) Clinical characteristics of pre-chemotherapy, post-chemotherapy, and both pre- and post-chemotherapy patients whose tumour tissue samples were used to construct the tissue microarray. (**b**) Cox regression analysis for overall survival as a function of age at diagnosis and residual disease after surgery in pre-chemotherapy patients, post-chemotherapy patients, and the entire cohort.

**Supplemental figure 6: Paired HGSOC tissue samples display senescence hallmarks following exposure to chemotherapy in patients.** (**a-f, n-o**) Matched pre-(left) and post-chemotherapy (right) tissue cores were stained in immunofluorescence for lamin B1 (**a**), Ki67 (**b**), IL6 (**c**), IL8 (**d**), vimentin (**e**), p16 (**f**), E-cadherin (**n**), cleaved caspase-3 (**o**), PML (**p**), MCM2 (**q**), and geminin (**r**). Nuclei are counterstained with DAPI (blue). Epithelial cells are counterstained with cytokeratin 7, 18 and 19 (fuscia). (**g-m, s-x**) Quantification of the MFI of lamin B1 (**g**), Ki67 (**h**), IL6 (**i**), IL8 (**j**), vimentin (**k**), p16 (**m**), E-cadherin (**p**), cleaved caspase-3 (**q**), PML (**p**), MCM2 (**q**), and geminin (**r**) in the epithelial (left) and stromal (right) tissue compartments of matched pre- and post-chemotherapy tissue cores. IL6, IL8, vimentin, E-cadherin, and cleaved caspase-3 MFIs were quantified in the entire epithelial or stromal compartment, whereas lamin B1, Ki67, p16, PML, MCM2, and geminin MFIs were quantified in the nucleus. The red data points correspond to the cores used in the representative images in (**a-f, n-r**).

**Supplemental figure 7: Senescence-associated biomarker levels in post-chemotherapy HGSOC tissues correlate with 5-year survival.** (**a-b**) Table summarizing the Kaplan-Meier log rank p-value comparing high and low pre- and post-chemotherapyexpression of different senescence markers, as well as expression independent of chemotherapy status, in the epithelial (**a**) and stromal (**b**) compartments in terms of 5-year survival. Whether higher or lower expression of the marker is advantageous for survival is indicated as well. Groups were separated into high and low expression based on cut-offs determined by ROC analyses.

**Supplemental figure 8: Pre- vs. post-chemotherapy status in patients with resectable tumour at the time of surgery predicts 5-year overall survival**. (**a**) Kaplan-Meier analysis comparing 5-year overall survival between pre- (primary debulking surgery followed by adjuvant chemotherapy) (n = 70) and post- (neoadjuvant chemotherapy followed by interval debulking surgery) (n = 72) patient groups.

**Supplemental figure 9: MFI correlations between duplicate cores.** (**a**) Pearson correlations of the MFIs of different markers between duplicate cores in the epithelial (**a**) and stromal (**b**) compartments in the full set of cores on the TMA.

**Supplemental table 1:** Summary of the characteristics of HGSOC primary cultures used for gene expression profiling: Origins, passages in culture and E2F-response cluster classification.

**Supplemental table 2:**
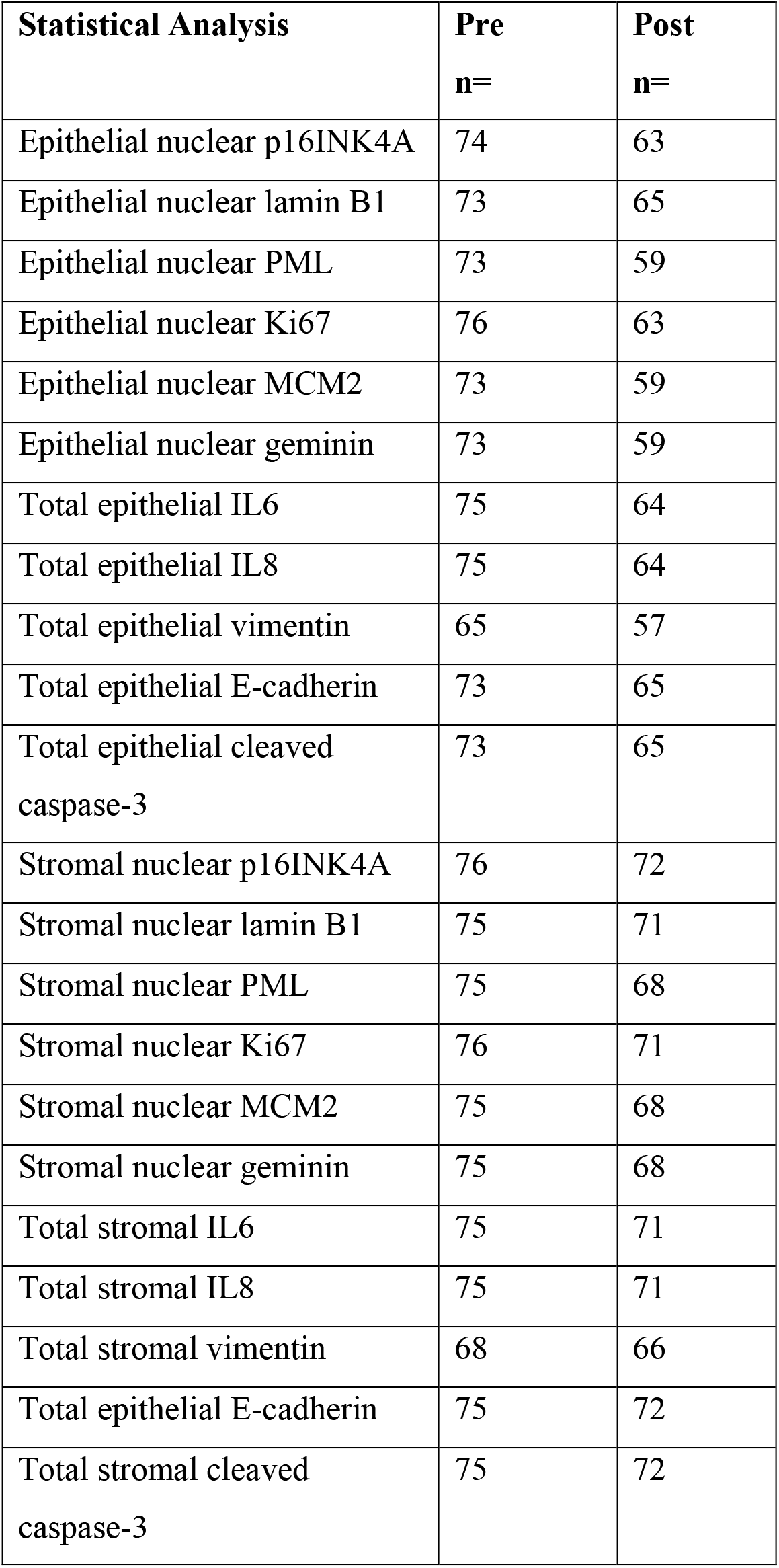
Breakdown of usable TMA cores per analysis performed: Exact number of cores in pre- and post-treatment groups that were included for statistical analysis.

**Supplemental table 3:** Summary of primary and secondary antibodies used in this study.

