## Supplemental figures for "A senescence-like state is beneficial for ovarian cancer treatment"

**Supplemental Figure 1: Workflow for HGSOC primary cell senescence transcriptome analysis.**

**a**

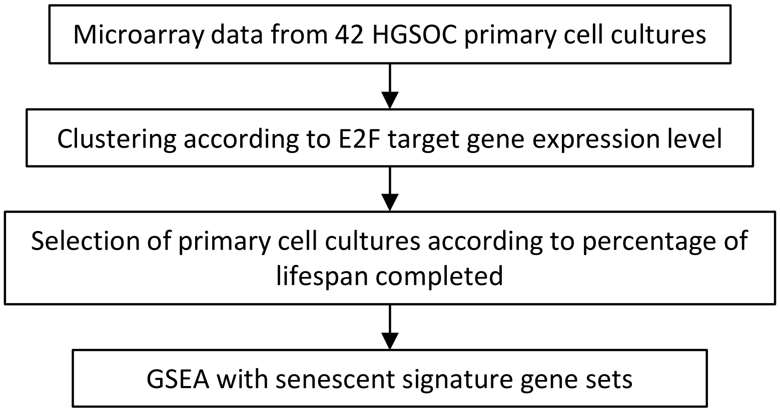


**b**

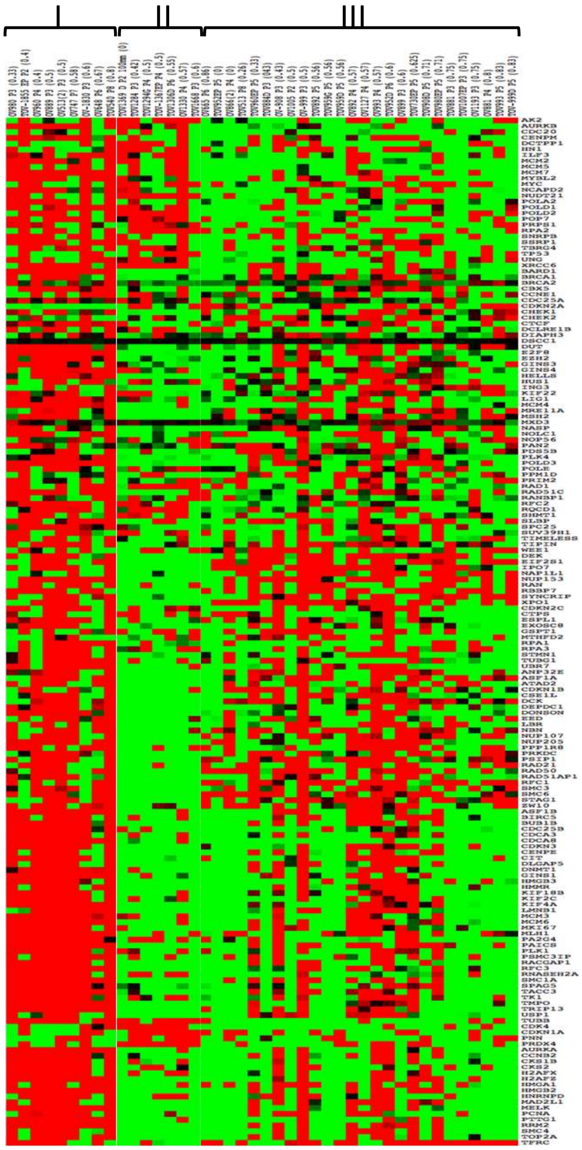


**c**

| Early passage primary cell cultures from Cluster I (≤0.5 of eventual maximal passage number), n=5 | Late passage primary cell cultures from Cluster III (≥0.75 of eventual maximal passage number), n=7 |
|---|---|
| OV980 P3 (0.33) | TOV881 P3 (0.75) |
| TOV1855 EP P2 (0.4) | TOV1007 EP P3 (0.75) |
| OV960 P4 (0.5) | OV1193 P3 (0.75) |
| OV889 P3 (0.5) | OV881 P4 (0.8) |
| OV513 P3 (0.5) | TOV993 P5 (0.83) |
| | TOV999D P5 (0.83) |
| | OV665 P6 (0.86) |

**d**

| 9 senescent signature gene sets are significantly enriched (p<0.01) in “Late” primary cell cultures (≥0.75 of eventual maximal passage number) | 6 senescent signature gene sets are significantly enriched (p<0.01) in “Early” primary cell cultures (≤0.50 of eventual maximal passage number) |
|---|---|
| Signature for Replicative Senescence_UP | Signature for Replicative Senescence_DN |
| Signature of IRIS at day 4 post-irradiation (genes shared among fibroblasts, melanocytes and keratinocytes)_UP | Signature of IRIS at day 4 post-irradiation (genes shared among fibroblasts, melanocytes and keratinocytes)_DN |
| Genes in the Fibroblast Signature of Senescence Independent of Stimulus_UP | Signature of IRIS at day 20 post-irradiation (genes shared among fibroblasts, melanocytes and keratinocytes)_DN |
| Signature for Ionizing-Radiation Induced Senescence_UP | Genes in the Fibroblast Signature of Senescence Independent of Stimulus_DN |
| Signature of IRIS at day 20 post-irradiation (genes shared among fibroblasts, melanocytes and keratinocytes)_UP | Signature of IRIS at day 10 post-irradiation (genes shared among fibroblasts, melanocytes and keratinocytes)_DN |
| Signature of Melanocytes_DN | Signature of Keratinocytes_DN |
| Signature of Astrocytes_UP | |
| Signature of Keratinocytes_UP | |
| Signature for Oncogene Induced Senescence_DN | |

**Supplemental Table 1: Characteristics of the primary cell cultures used in this work.**

| HGSC Patient ID | Origin of tumor cells | Primary cells | Passage of RNA extraction | Maximal number of passages | Relative Lifespan ratio | E2F Cluster category |
|---|---|---|---|---|---|---|
| 513 | ovary | TOV513 | 8 | 31 | 0.2580645 | III |
| | ascites | OV513(2) | 3 | 6 | 0.5 | I |
| 540 | ovary | TOV540 | 8 | 10 | 0.8 | I |
| 648 | ascites | OV648 | 6 | 9 | 0.6666667 | I |
| 665 | ascites | OV665 | 6 | 7 | 0.8571429 | III |
| 730 | ovary | TOV730EP | 5 | 8 | 0.625 | III |
| 747 | ascites | OV747 | 7 | 12 | 0.5833333 | I |
| 866 | ascites | OV866(2) | 4 | 100 | 0.04 | III |
| 881 | ovary | TOV881 | 3 | 4 | 0.75 | III |
| | ascites | OV881 | 4 | 5 | 0.8 | III |
| 884 | ovary | TOV884D | 3 | 7 | 0.4285714 | III |
| 889 | ascites | OV889 | 3 | 6 | 0.5 | I |
| 892 | ovary | TOV892 | 5 | 9 | 0.5555556 | III |
| | ascites | OV892 | 4 | 7 | 0.5714286 | III |
| 899 | ascites | OV899 | 3 | 5 | 0.6 | III |
| 908 | ascites | OV-908 | 3 | 7 | 0.4285714 | III |
| | ovary | TOV908D | 5 | 7 | 0.7142857 | III |
| 952 | omentum | TOV952EP | 5 | 100 | 0.05 | III |
| | ovary | TOV952D | 6 | 10 | 0.6 | III |
| 959 | ovary | TOV959G | 5 | 9 | 0.5555556 | III |
| | ovary | TOV959D | 5 | 9 | 0.5555556 | III |
| 960 | omentum | TOV960EP | 5 | 15 | 0.3333333 | III |
| | ascites | OV960 | 4 | 10 | 0.4 | I |
| 980 | ascites | OV980 | 3 | 9 | 0.3333333 | I |
| | omentum | TOV980EP | 5 | 7 | 0.7142857 | III |
| 993 | ascites | OV993 | 4 | 7 | 0.5714286 | III |
| | ovary | TOV993 | 5 | 6 | 0.8333333 | III |
| 999 | ascites | OV-999 | 3 | 6 | 0.5 | III |
| | ovary | TOV-999D | 5 | 6 | 0.8333333 | III |
| 1005 | ascites | OV1005 | 2 | 4 | 0.5 | III |
| 1007 | omentum | TOV1007EP | 3 | 4 | 0.75 | III |
| 1127 | ascites | OV1127 | 4 | 7 | 0.5714286 | III |
| 1193 | ascites | OV1193 | 3 | 4 | 0.75 | III |
| 1284 | ovary | TOV1284 | 3 | 7 | 0.4285714 | II |
| 1294 | ovary | TOV1294G | 4 | 8 | 0.5 | II |
| 1306 | ovary | TOV1306D | 6 | 11 | 0.5454545 | II |
| 1330 | ascites | OV1330 | 4 | 7 | 0.5714286 | II |
| 1367 | omentum | TOV-1367EP | 4 | 8 | 0.5 | II |
| 1369 | ovary | TOV1369D | 2 | 100 | 0.02 | II |
| 1668 | ovary | TOV1668 | 3 | 5 | 0.6 | II |
| 1830 | ascites | OV1830 | 3 | 5 | 0.6 | I |
| 1855 | omentum | TOV-1855EP | 2 | 5 | 0.4 | I |

**Supplemental Table 2: Breakdown of usable TMA cores per analysis performed.**

| Statistical Analysis | Pre n= | Post n= |
|---|---|---|
| Epithelial nuclear p16INK4A | 74 | 63 |
| Epithelial nuclear lamin B1 | 73 | 65 |
| Epithelial nuclear PML | 73 | 59 |
| Epithelial nuclear Ki67 | 76 | 63 |
| Epithelial nuclear MCM2 | 73 | 59 |
| Epithelial nuclear geminin | 73 | 59 |
| Total epithelial IL6 | 75 | 64 |
| Total epithelial IL8 | 75 | 64 |
| Total epithelial vimentin | 65 | 57 |
| Total epithelial E-cadherin | 73 | 65 |
| Total epithelial cleaved caspase-3 | 73 | 65 |
| Stromal nuclear p16INK4A | 76 | 72 |
| Stromal nuclear lamin B1 | 75 | 71 |
| Stromal nuclear PML | 75 | 68 |
| Stromal nuclear Ki67 | 76 | 71 |
| Stromal nuclear MCM2 | 75 | 68 |
| Stromal nuclear geminin | 75 | 68 |
| Total stromal IL6 | 75 | 71 |
| Total stromal IL8 | 75 | 71 |
| Total stromal vimentin | 68 | 66 |
| Total epithelial E-cadherin | 75 | 72 |
| Total stromal cleaved caspase-3 | 75 | 72 |

**Supplemental Table 3: Summary of primary and secondary antibodies used in this study.**

| Antibody (clone, if monoclonal) | Species | Manufacturer (catalog number) | Dilution for primary cell IF | Dilution for TMA IF |
|---|---|---|---|---|
| **Primary antibodies** | | | | |
| p16 (JC8) | Mouse | Santa Cruz (sc-56330) | 1:100 or 1:300 | 1:500 |
| Ki-67 (SP6) | Rabbit | ThermoFisher Scientific (RM-9106-S) | - | 1:500 |
| Lamin B1 | Rabbit | Abcam (ab16048) | 1:1000 | 1:500 |
| Cleaved caspase-3 | Rabbit | Cell Signaling (6991) | - | 1:500 |
| IL8 (6217) | Mouse | R&D Systems (MAB208) | - | 1:50 |
| IL6 | Rabbit | Abcam (ab6672) | - | 1:500 |
| Vimentin (V9) | Mouse | Sigma-Aldrich (V6630) | - | 1:100 |
| E-cadherin (G10) | Mouse | Santa Cruz (sc-8426) | - | 1:50 |
| 53BP1 | Rabbit | Novus Biologicals (NB100-304) | 1:2000 | - |
| PML | Goat | Santa Cruz (sc-9862) | 1:500 | 1:500 |
| MCM2 (1E7) | Mouse | Cell Signaling (12079) | - | 1:200 |
| Geminin | Rabbit | Proteintech (10802-1-AP) | - | 1:5000 |
| **Epithelial mask primary antibodies** | | | | |
| Cytokeratin 7 (OV-TL 12/30) | Mouse | ThermoFisher Scientific (MS-1352-P) | - | 1:200 |
| Cytokeratin 18 (DC-10) | Mouse | Santa-Cruz (sc-6259) | - | 1:200 |
| Keratin 19 (A53-B/A2.26) | Mouse | ThermoFisher Scientific (MS-198-P) | - | 1:200 |
| **Secondary antibodies** | | | | |
| Alexa Fluor 488 anti-mouse | Donkey | Life Technologies (A21202) | 1:800 | - |
| Alexa Fluor 568 anti-mouse | Donkey | Life Technologies (A10037) | - | 1:250 |
| Alexa Fluor 568 anti-rabbit | Donkey | Life Technologies (A10042) | - | 1:250 |
| Alexa-Fluor 647 anti-rabbit | Donkey | Life Technologies (A31573) | 1:800 | 1:250 |
| Alexa-Fluor 647 anti-mouse | Donkey | Life Technologies (A31571) | 1:800 | 1:250 |
| Alexa-Fluor 750 anti-mouse | Goat | Life Technologies (A21037) | - | 1:250 |
| Cy3 anti-goat | Donkey | Jackson ImmunoResearch (705-165-147) | 1:800 | - |

**Supplemental Figure 3: HGSOC primary cultures undergo TIS in response to DNA damage and chemotherapy.**

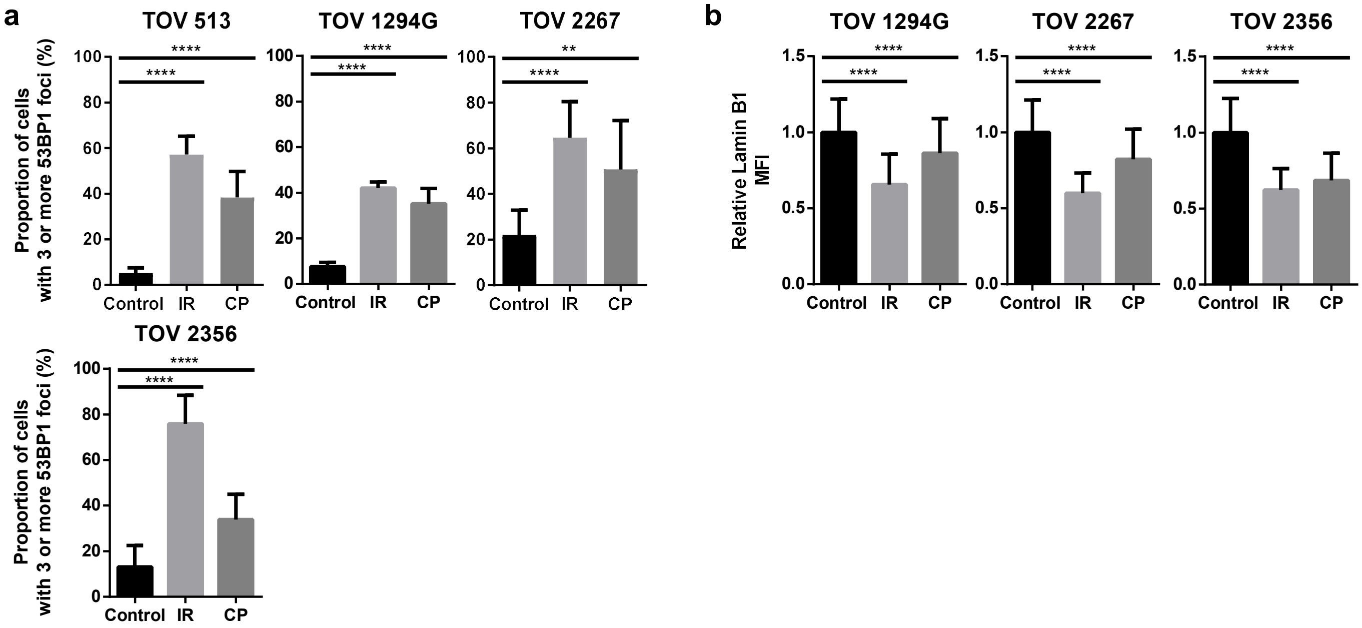

**Supplemental Figure 4: HGSOC TIS renders cells sensitive to ABT-263.**

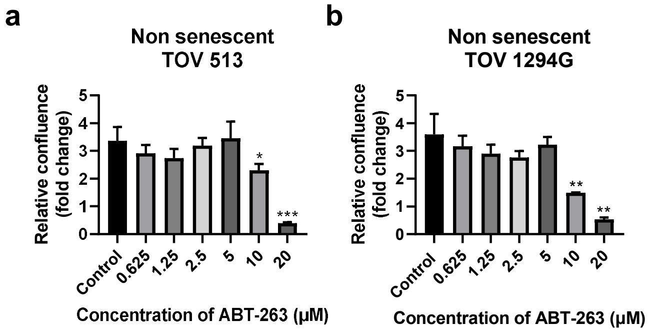

**Supplemental Figure 5: Clinical characteristics of the patients in the pre- and post-chemotherapy HGSOC cohort used to construct the tissue microarray.**

**a**

| | Total n=148 | Pre-chemo n=76 | Post-chemo n=72 |
|---|---|---|---|
| | | Mean (SD) | |
| Age at Diagnosis (years) | 60.21 (9.61) | 60.54 (9.70) | 59.86 (9.56) |
| | | Median (SD) | |
| Follow up (months) | 32.00 (35.286) | 46.50 (40.03) | 25.50 (27.12) |
| | | n (%) | |
| Disease-specific vital status within 5 years follow-up | | | |
| Alive | 65 (43.9 %) | 35 (46.1 %) | 30 (41.7 %) |
| Deceased | 83 (56.1 %) | 41 (53.9 %) | 42 (58.3 %) |
| FIGO stage at presentation | | | |
| 1 | 2 (1.4 %) | 2 (2.6 %) | 0 (0.0 %) |
| 2 | 7 (4.7%) | 6 (7.9 %) | 1 (1.4 %) |
| 3 | 118 (79.7 %) | 60 (78.9 %) | 58 (80.6 %) |
| 4 | 19 (12.8 %) | 8 (10.5 %) | 11 (15.3 %) |
| Unknown | 2 (1.4 %) | 0 (0.0 %) | 2 (2.8 %) |
| Residual Disease After Surgery | | | |
| None | 33 (22.3 %) | 12 (15.8 %) | 21 (29.2 %) |
| 1 cm or less | 37 (25.0 %) | 19 (25.0 %) | 28 (25.0 %) |
| 1-2 cm | 19 (12.8 %) | 11 (14.5 %) | 8 (11.1 %) |
| 2 cm and greater | 33 (22.3 %) | 26 (34.2 %) | 7 (9.7 %) |
| Milliary | 7 (4.7 %) | 3 (3.9 %) | 4 (5.6 %) |
| Unknown | 19 (12.8 %) | 5 (6.6 %) | 14 (19.4 %) |

**b**

| | Overall 5-year survival | | | | | | | | |
|---|---|---|---|---|---|---|---|---|---|
| | Total | | | Pre-chemo | | | Post-chemo | | |
| Variable | HR[a] | 95 % CI[b] | p-value | HR[a] | 95 % CI[b] | p-value | HR[a] | 95 % CI[b] | p-value |
| Age at diagnosis | 1.006 | 0.982-1.031 | 0.613 | 1.009 | 0.971-1.048 | 0.649 | 1.008 | 0.977-1.039 | 0.620 |
| Residual disease after surgery | | | **0.0475** | | | 0.115 | | | **0.0119** |
| 1 cm or less vs none | 1.351 | 0.637-2.865 | 0.433 | 2.495 | 0.518-12.02 | 0.254 | 1.111 | 0.458-2.694 | 0.816 |
| 1-2 cm vs none | 2.311 | 1.035-5.161 | **0.0410** | 3.863 | 0.779-19.16 | 0.0981 | 2.837 | 1.051-7.655 | **0.0395** |
| 2 cm and greater vs none | 2.558 | 1.224-5.348 | **0.0125** | 5.725 | 1.314-24.95 | **0.0202** | 2.381 | 0.730-7.769 | 0.151 |
| Miliary vs none | 3.036 | 0.965-9.546 | 0.0575 | 4.416 | 0.621-31.39 | 0.138 | 12.635 | 2.405-66.37 | **0.00273** |

[a]Hazard ratio (HR) estimated from Cox proportional hazard regression model. [b]Confidence interval of the estimated HR.

**Supplemental Figure 6: HGSOC tissues display senescence hallmarks following exposure to chemotherapy in patients.**

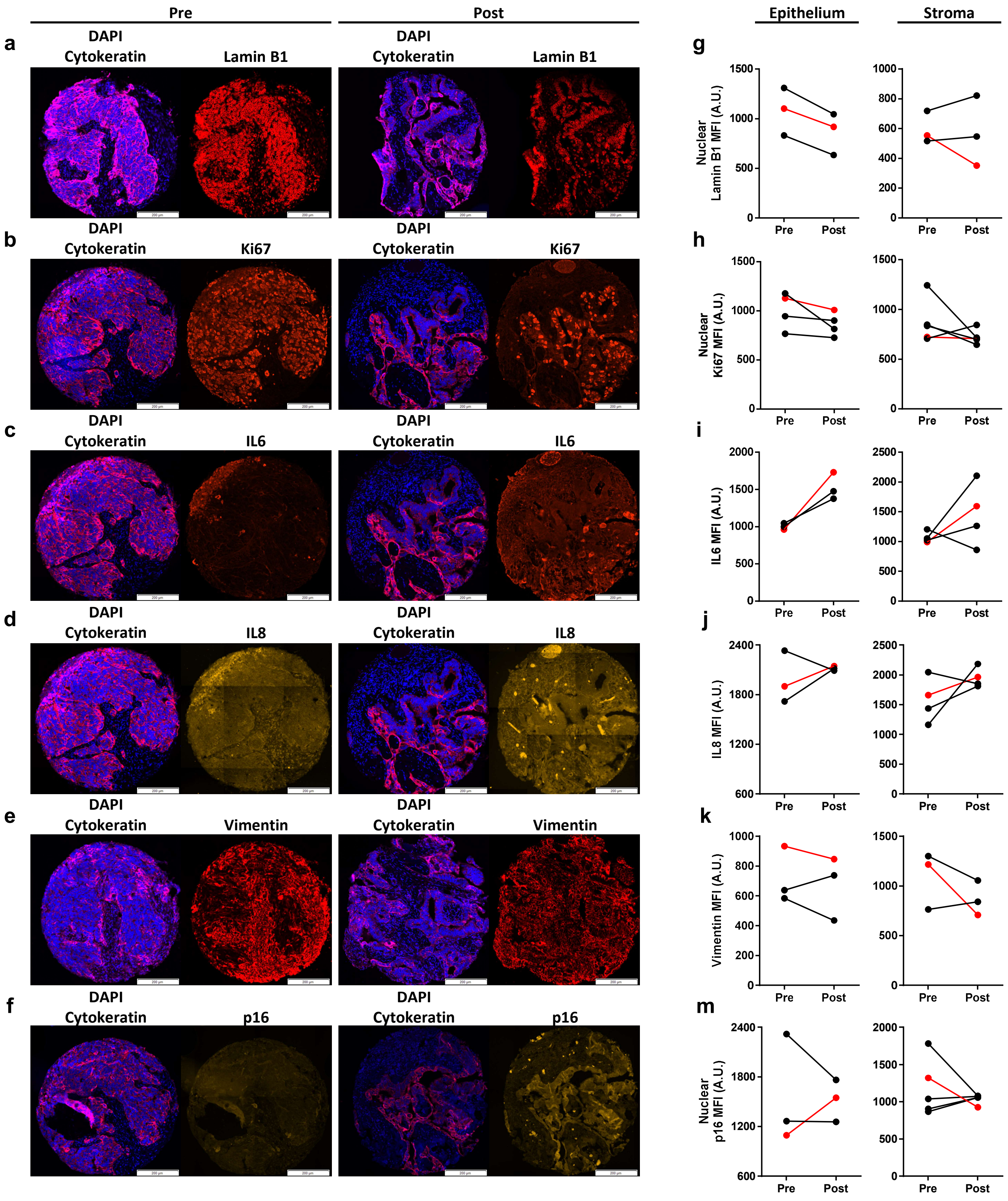

**Supplemental Figure 6 (cont'd): HGSOC tissues display senescence hallmarks following exposure to chemotherapy in patients.**

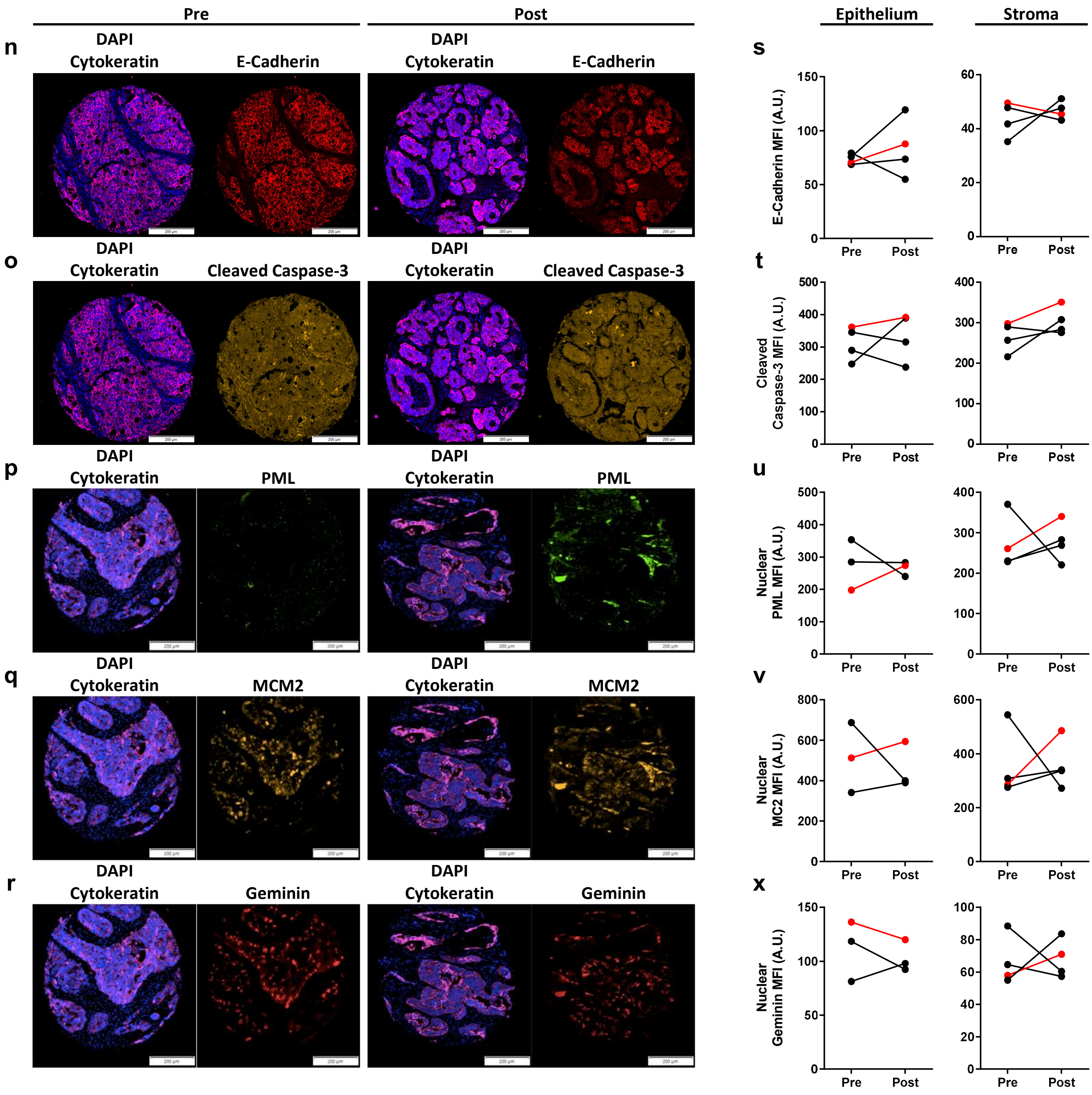

**Supplemental figure 7: Senescence-associated marker levels in post-chemotherapy HGSOC tissues correlate with 5-year survival.**

**a**

| | Epithelium | | | | | |
|---|---|---|---|---|---|---|
| | Pre | | Post | | All | |
| **Marker** | **Survival Advantage if Marker** | **Kaplan-Meier Log Rank p-value** | **Survival Advantage if Marker** | **Kaplan-Meier Log Rank p-value** | **Survival Advantage if Marker** | **Kaplan-Meier Log Rank p-value** |
| **Nuclear p16** | High | **0.019** | High | **0.017** | High | **0.002** |
| **Nuclear Lamin B1** | High | 0.159 | Low | **0.045** | Low | **0.009** |
| **Nuclear PML** | High | 0.275 | Low | 0.224 | High | 0.312 |
| **Nuclear Ki67** | High | 0.248 | Low | 0.129 | High | 0.294 |
| **Nuclear MCM2** | Low | 0.056 | Low | **0.015** | Low | **0.013** |
| **Nuclear Geminin** | High | 0.057 | Low | 0.056 | High | 0.203 |
| **IL6** | High | 0.058 | Low | 0.305 | High | 0.148 |
| **IL8** | Low | **0.027** | High | 0.285 | Low | 0.857 |
| **Cleaved Caspase-3** | High | 0.196 | Low | 0.562 | High | 0.452 |
| **Vimentin** | Low | **0.011** | Low | 0.287 | Low | **0.009** |
| **E-Cadherin** | High | 0.091 | Low | 0.951 | Low | 0.442 |
| **$G_0$ ($Ki67^-/MCM2^-$)** | Low | 0.178 | Low | 0.390 | Low | 0.097 |
| **$G_1$ ($Ki67^-/MCM2^+$)** | Low | **0.022** | Low | 0.596 | Low | 0.895 |
| **S ($Ki67^+/MCM2^+$)** | High | 0.091 | Low | 0.407 | High | 0.109 |
| **G2/M ($Ki67^+/MCM2^-$)** | High | 0.240 | High | 0.101 | High | 0.140 |
| **$G_0$ ($Geminin^-/MCM2^-$)** | Low | 0.230 | High | 0.053 | High | 0.133 |
| **$G_1$ ($Geminin^-/MCM2^+$)** | High | **0.013** | High | 0.080 | High | **0.008** |
| **S ($Geminin^+/MCM2^+$)** | High | **0.032** | Low | 0.128 | High | 0.398 |
| **G2/M ($Geminin^+/MCM2^-$)** | Low | **0.015** | Low | **0.039** | Low | **0.010** |

**b**

| | Stroma | | | | | |
|---|---|---|---|---|---|---|
| | Pre | | Post | | All | |
| **Marker** | **Survival Advantage if Marker** | **Kaplan-Meier Log Rank p-value** | **Survival Advantage if Marker** | **Kaplan-Meier Log Rank p-value** | **Survival Advantage if Marker** | **Kaplan-Meier Log Rank p-value** |
| **Nuclear p16** | High | 0.413 | Low | 0.789 | High | 0.332 |
| **Nuclear Lamin B1** | High | **<0.001** | Low | 0.428 | High | **0.022** |
| **Nuclear PML** | High | 0.087 | Low | 0.340 | Low | 0.072 |
| **Nuclear Ki67** | High | 0.100 | High | 0.132 | High | **0.014** |
| **Nuclear MCM2** | High | 0.528 | Low | 0.169 | High | 0.694 |
| **Nuclear Geminin** | High | 0.080 | Low | 0.239 | High | 0.120 |
| **IL6** | High | 0.230 | High | 0.103 | High | **0.007** |
| **IL8** | High | 0.071 | High | 0.062 | High | **0.021** |
| **Cleaved Caspase-3** | High | 0.624 | High | 0.968 | High | 0.903 |
| **Vimentin** | High | 0.415 | Low | **0.001** | Low | **0.001** |
| **E-Cadherin** | Low | 0.703 | High | **0.042** | High | 0.358 |
| **$G_0$ ($Ki67^-/MCM2^-$)** | Low | **0.001** | Low | 0.538 | Low | 0.110 |
| **$G_1$ ($Ki67^-/MCM2^+$)** | Low | 0.664 | High | 0.248 | High | 0.376 |
| **S ($Ki67^+/MCM2^+$)** | High | **0.021** | High | 0.090 | High | 0.175 |
| **G2/M ($Ki67^+/MCM2^-$)** | High | 0.337 | High | 0.245 | High | 0.114 |
| **$G_0$ ($Geminin^-/MCM2^-$)** | Low | 0.083 | High | 0.240 | Low | **0.035** |
| **$G_1$ ($Geminin^-/MCM2^+$)** | High | **0.017** | High | **0.020** | High | **<0.001** |
| **S ($Geminin^+/MCM2^+$)** | High | 0.051 | Low | 0.200 | High | 0.105 |
| **G2/M ($Geminin^+/MCM2^-$)** | High | 0.164 | Low | 0.500 | High | 0.159 |

**Supplemental figure 8: Pre- vs. post-chemotherapy status in patients with resectable tumour at the time of surgery predicts 5-year overall survival.**

a

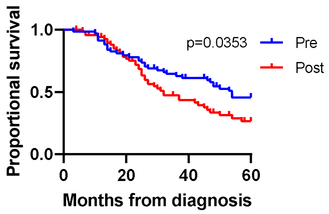

### Supplemental Figure 9 : MFI correlations between duplicate cores.

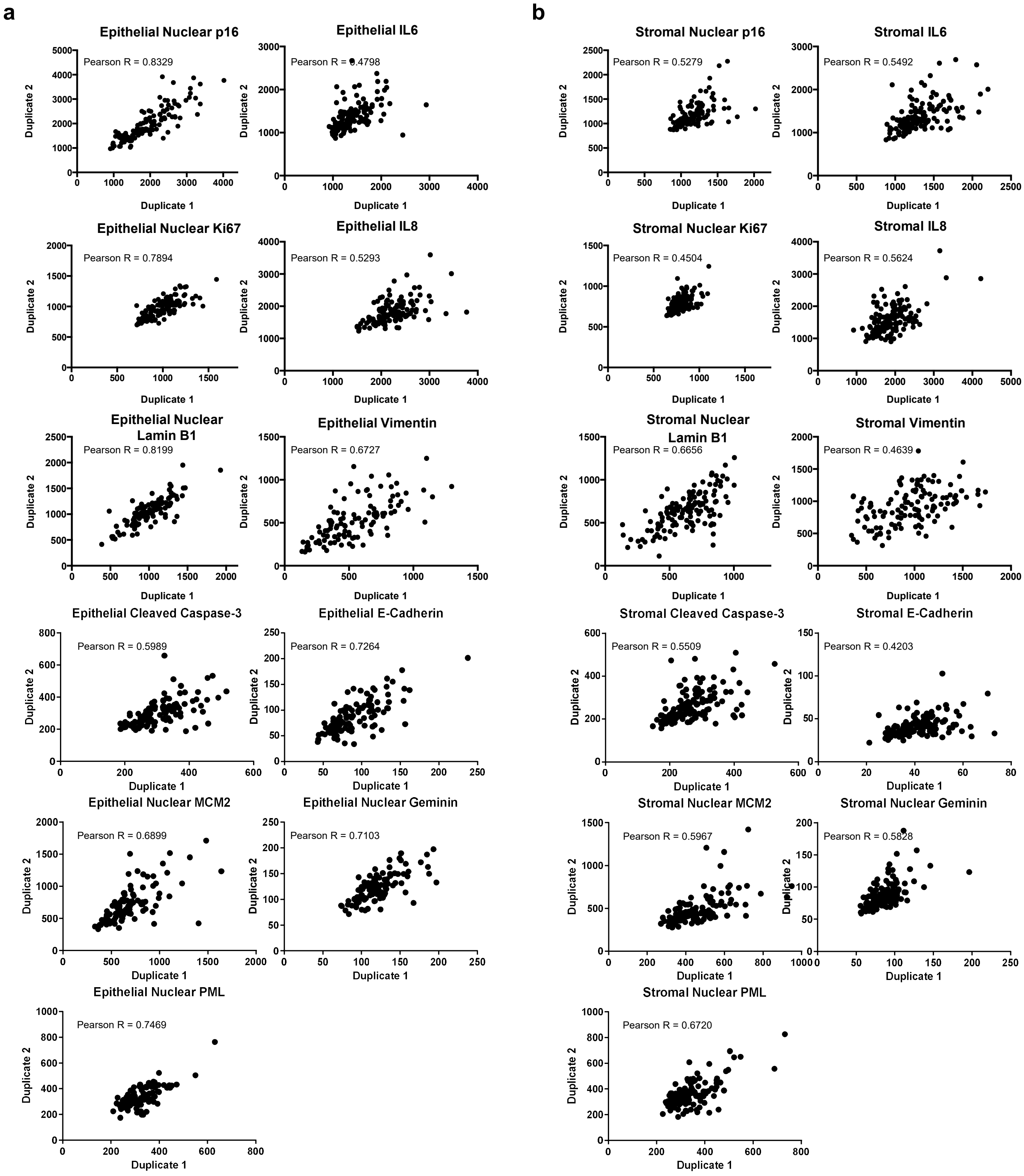
